# “The *B. subtilis* translesion polymerase Pol Y1 is not strongly recruited to sites of replication upon different types of DNA damage”

**DOI:** 10.64898/2026.04.02.716108

**Authors:** Sophia R. Martinez-Whitman, Chloe M. Santana, Alyssa P. Campbell, Denholm T. Feldman, Isaac E.Z. Jabaley, Luke G. O’Neal, McKayla E. Marrin, Elizabeth S. Thrall

## Abstract

One challenge to DNA replication is the presence of unrepaired damage on the template strand, which can stall the replication machinery. This stall can be resolved by the translesion synthesis (TLS) pathway, in which specialized translesion polymerases are recruited to copy damaged DNA. Because TLS polymerases are error-prone, their activity is regulated at multiple levels to minimize unnecessary mutagenesis. Although the molecular mechanisms of bacterial TLS have been extensively studied in *Escherichia coli*, less is known about this pathway in other species. In *E. coli*, the TLS polymerase Pol IV is minimally enriched at replication forks in the absence of DNA damage but is strongly recruited upon replication stalling, enabling TLS while minimizing mutagenesis. However, we recently showed that the *Bacillus subtilis* TLS polymerase Pol Y1, the homolog of Pol IV, is moderately enriched near replication sites even during normal growth and is not further enriched upon treatment with the DNA damaging agent 4-nitroquinoline 1-oxide (4-NQO). It is unknown whether this behavior is unique to 4-NQO or general to other types of DNA damage. In this study, we investigate the effects of four different DNA damaging agents (ultraviolet light, methyl methanesulfonate, nitrofurazone, and mitomycin C) in *B. subtilis*. We first characterize the contributions of the two TLS polymerases, Pol Y1 and Pol Y2, to DNA damage survival and damage-induced mutagenesis after treatment with these agents. We then use single-molecule fluorescence microscopy to measure the localization and dynamics of individual Pol Y1 molecules in live *B. subtilis* cells. We find that Pol Y1 and Pol Y2 have differing effects on survival and mutagenesis, but that under no circumstances is Pol Y1 strongly recruited to sites of replication upon DNA damage. This study broadens our understanding of TLS in *B. subtilis*, indicating that there are notable differences in TLS mechanisms across bacteria.

## Introduction

Unrepaired DNA damage can stall the replication machinery, a multi-protein complex known as the replisome, ultimately leading to cell death. Translesion synthesis (TLS) is a DNA damage tolerance pathway that promotes cell survival by alleviating this stall.(1–3) In this process, specialized translesion polymerases, generally members of the error-prone Y family, are recruited to copy damaged DNA, allowing replication to continue past blocking lesions. Although TLS promotes cell survival, TLS polymerases are generally low-fidelity, and therefore TLS is a mutagenic process. Thus, cellular regulation of TLS polymerases must strike a balance between minimizing their activity during normal unstressed conditions while also allowing them access to the DNA template when replication is stalled.

In bacteria, TLS has primarily been studied in the model gram-negative species *Escherichia coli*. In *E. coli*, it is clear that TLS polymerases are regulated at multiple levels to maintain genome stability while also allowing them access to stalled replication forks.(2) In particular, the TLS polymerase Pol IV is regulated transcriptionally via the SOS DNA damage response, which increases its cellular copy number approximately 10-fold after SOS induction. In addition to this transcriptional regular, recent single-molecule imaging studies have also revealed a spatial component to Pol IV regulation. Even when Pol IV copy number is held constant, Pol IV is not strongly enriched at sites of replication during normal replication but is strongly enriched upon DNA damage or other replication perturbations, even for non-cognate DNA lesions that it cannot bypass.(4–6) These mechanisms ensure that Pol IV can gain access to the DNA template upon replication stalling, while limiting its access during unperturbed replication.

Although the molecular mechanisms of TLS have been elucidated in great detail in *E. coli*, much less is known about TLS in other bacterial species. The model gram-positive bacterium *Bacillus subtilis* is an attractive species to serve as an alternative model for bacterial TLS, given its evolutionary distance from *E. coli*. *B. subtilis* has two TLS polymerases, Pol Y1 and Pol Y2, which are the homologs of *E. coli* Pol IV and Pol V, respectively.(7–9) A few studies have started to reveal similarities and differences in the contributions of Pol Y1 and Pol Y2 to DNA damage survival and mutagenesis relative to their *E. coli* homologs. Like Pol IV, Pol Y1 promotes survival to the drug 4-nitroquinoline 1-oxide (4-NQO).(10,11) However, unlike Pol IV, Pol Y1 promotes survival upon ultraviolet C (UV-C) treatment, whereas Pol Y2 deletion has no effect.(10,12,13) In contrast, *E. coli* Pol V mediates survival to UV damage.(14,15) Both Pol Y1 and Pol Y2 were shown to play a role in the tolerance of hexavalent chromium (Cr(VI))-mediated damage.(16) One study investigated DNA damage tolerance and mutagenesis in sporulating cells, rather than vegetatively growing cells, finding that both Pol Y1 and Pol Y2 had an effect on survival and mutagenesis for damaging agents like UV-C light, mitomycin C (MMC), and *tert*-butyl hydroperoxide.(12)

In addition to differences in the functional roles of Pol Y1 and Pol Y2, it is becoming clear that there are differences in their cellular regulation. Unlike the *E. coli* TLS polymerases and Pol Y2, Pol Y1 is not a member of the SOS regulon.(8,17) Further, although Pol V is composed of two different subunits, the catalytic subunit UmuC and the accessory subunit UmuD,(2) Pol Y2 is currently thought to function as a single gene product.(8) There is also evidence that the spatial regulation of Pol Y1 differs from that of Pol IV. In a previous study, we found that Pol Y1 is moderately enriched at replication sites during normal growth, but that it is not further enriched upon 4-NQO treatment,(11) in contrast to Pol IV.(4,6) Similarly, we found no changes in Pol Y1 mobility upon 4-NQO treatment,(11) whereas DNA damage was shown to lead to increased Pol IV binding.(4) However, it is not known whether this lack of Pol Y1 recruitment is unique to 4-NQO or whether it is a general phenomenon.

In this study, we determine the contributions of Pol Y1 and Pol Y2 to DNA damage survival and mutagenesis in response to treatment with four different DNA damaging agents: UV light, methyl methanesulfonate (MMS), nitrofurazone (NFZ), and MMC. In matched experiments, we image single Pol Y1 molecules in live *B. subtilis* cells and measure their cellular localization and mobility. Although Pol Y1 and Pol Y2 deletions have differing effects on cell survival and mutagenesis upon different types of DNA damage, we find relatively modest changes in Pol Y1 mobility and in the enrichment of Pol Y1 at replication sites, even for cognate DNA damaging agents. These results indicate that the lack of Pol Y1 recruitment observed previously for 4-NQO is consistent for other types of DNA damage and reveal further differences in the mechanisms of TLS in *E. coli* and *B. subtilis*.

## Results

### Pol Y1 deletion has differing effects on cell survival, whereas Pol Y2 deletion has no effect

To survey the contributions of Pol Y1 and Pol Y2 to cell survival upon DNA damage, we decided to test treatment with four different DNA damaging agents that generate a range of different lesions. First, we tested the effect of UV irradiation, which generates highly blocking DNA lesions like cyclobutane pyrimidine dimers and 6–4 photoproducts.(18) In *E. coli*, Pol IV does not contribute to UV survival; instead, Pol V performs TLS past UV-induced lesions.(19,14,15) In *B. subtilis*, previous work by our lab(13) and others(7,10) has shown that only Pol Y1 promotes survival to UV damage, in contrast to *E. coli*. We assayed survival of cells to 0, 10, 20, and 40 J/m^2^ doses of 254 nm UV-C light (Figure 1A) and compared the WT strain to strains bearing knockouts of Pol Y1 (Δ*yqjH*) and Pol Y2 (Δ*yqjW*) individually or in combination (Δ*yqjH* Δ*yqjW*). Consistent with prior reports,(7,10) we found that deletion of Pol Y1 sensitized cells to UV (by roughly 10-fold at the highest dose), whereas deletion of Pol Y2 had no effect. The sensitivity of the double knockout alone was comparable to that of the single Pol Y1 knockout, indicating that Pol Y2 cannot substitute for Pol Y1 to bypass UV lesions.

**Figure 1.**
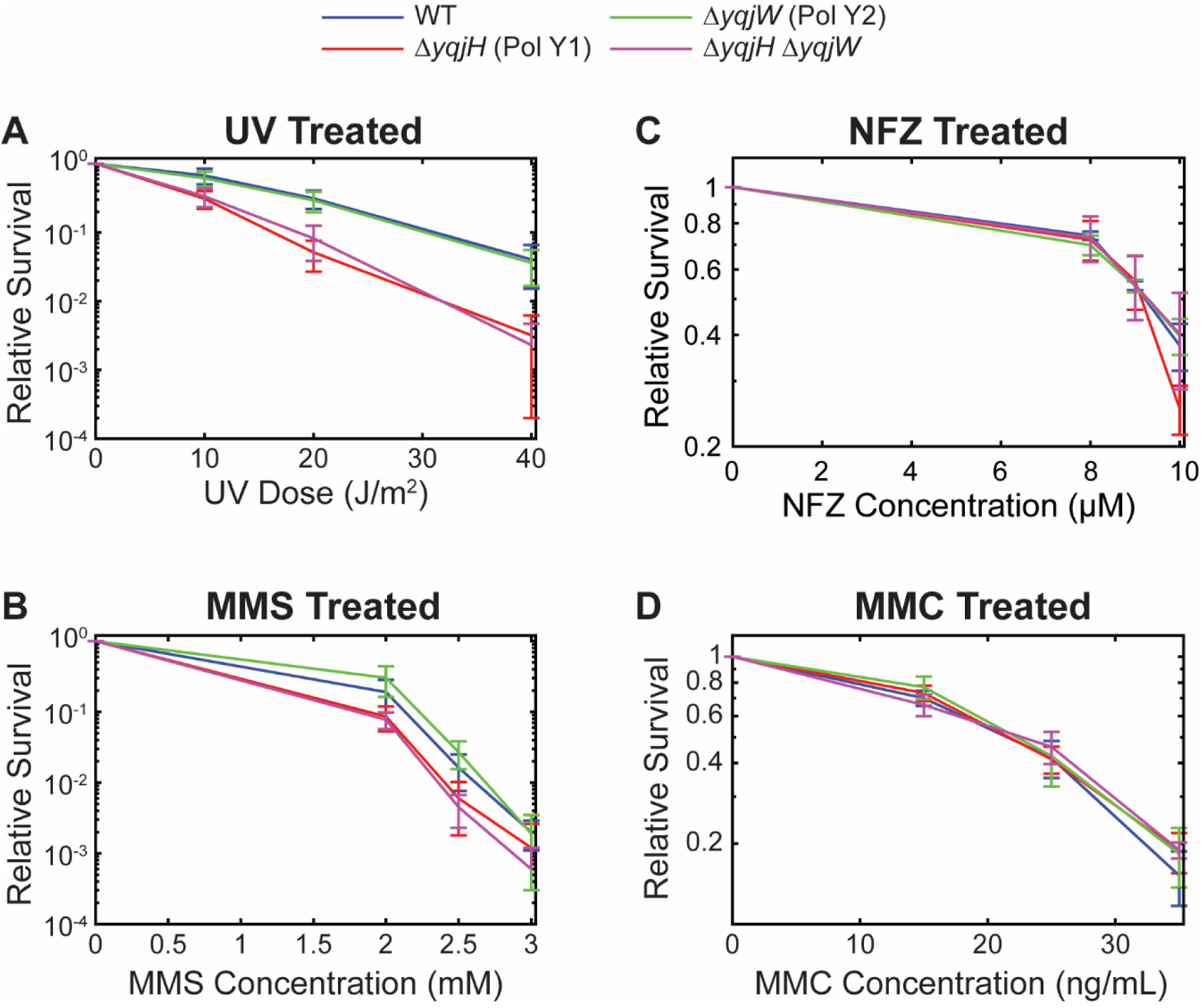
Survival of *B. subtilis* strains in the presence of different types of DNA damage. Relative survival rates for WT, ΔPol Y1 knockout, ΔPol Y2 knockout, and ΔPol Y1 ΔPol Y2 double knockout strains after treatment with different doses of (A) 254 nm UV light, (B) MMS, (C) NFZ, and (D) MMC. Error bars show standard deviation of at least three replicates.

Next, we tested the effect of treatment with the alkylating agent MMS, which generates various DNA lesions, including the replication-blocking *N*^3^-methyladenine.(20,21) In *E. coli*, Pol IV bypasses MMS lesions and promotes cell survival, whereas Pol V does not contribute to tolerance.(22) However, no prior work has investigated the effect of Pol Y1 and Pol Y2 on MMS survival or mutagenesis in *B. subtilis*. We assayed survival of cells plated on LB agar plates containing MMS at concentrations of 0, 2, 2.5, and 3 mM (Figure 1B). As for UV treatment, we found that deletion of Pol Y1 sensitized cells to MMS, albeit more modestly (approximately two-to three-fold), whereas deletion of Pol Y2 had no effect. The sensitivity of the double knockout was comparable to that of Pol Y1 alone, again indicating that Pol Y2 cannot substitute for Pol Y1 to bypass MMS lesions.

The drug NFZ generates DNA strand breaks as well as small minor groove lesions, like *N*^2^ adducts of guanine,(23,24) which are efficiently bypassed by *E. coli* Pol IV, whereas Pol V does not promote survival.(15,25) Although NFZ is a well-studied cognate damaging agent for Pol IV, no prior work has investigated the effect of Pol Y1 and Pol Y2 on NFZ survival in *B. subtilis*. Therefore, we assayed survival of cells plated on LB agar plates containing NFZ at concentrations of 0, 8, 9, and 10 µM (Figure 1C). The survival rate was relatively high (∼ 35 – 40%) at the highest concentration and dropped precipitously above 10 µM (data not shown). However, we were unable to obtain consistent assay results at higher NFZ concentrations. Within this limited concentration range, there was no obvious effect of Pol Y1 or Pol Y2 deletion, either individually or in combination, with the possible exception of a slight decrease in survival for the Pol Y1 single knockout, but not the double knockout. Thus, in contrast to the behavior of its *E. coli* homolog, Pol IV does not appear to confer survival to NFZ damage, at least in the regime of relatively high survival rates.

Finally, we tested the effect of Pol Y1 and Pol Y2 deletion on survival upon treatment with the drug MMC. MMC is an alkylating agent that generates a wide range of different types of lesions, including both interstrand crosslinks (ICLs) and intrastrand crosslinks as well as *N*^2^ and *N*^7^ monoadducts of guanine.(26–29) To our knowledge, there are no reports on the roles of *E. coli* Pol IV and Pol V in tolerance of MMC-induced damage. In *B. subtilis*, no studies have tested the effect of Pol Y1 and Pol Y2 on survival or mutagenesis in vegetative cells, although both contribute to survival in sporulating cells.(12) We assayed survival of cells to 0, 15, 25, and 35 ng/mL concentrations of MMC in solid media (Figure 1D). At the highest of these concentrations, the relative survival of the WT strain was still relatively high, at approximately 0.15. We also tested a higher concentration of 45 ng/mL; however, at this concentration, survival dropped precipitously and the colonies became translucent and challenging to count. Thus, we did not use concentrations greater than 35 ng/mL. Over the range of concentrations, deletion of Pol Y1, Pol Y2, or both had no effect on MMC survival, in contrast to a previous report in sporulating cells.(12) These results suggest that Pol Y1 and Pol Y2 may play different roles during sporulation and vegetative growth.

### Pol Y1 and Pol Y2 both contribute to damage-induced mutagenesis under different conditions

In addition to surveying the roles of Pol Y1 and Pol Y2 in damage tolerance, we also characterized their effect on damage-induced mutagenesis upon treatment with the same set of damaging agents. We first explored UV-induced mutagenesis, which in *E. coli* is mediated by Pol V.(30) Prior results on UV-induced mutagenesis in *B. subtilis* were not in agreement. One study found that both Pol Y1 and Pol Y2 contribute, with Pol Y1 having the greater impact,(7) whereas another study reported that Pol Y2 alone was responsible for UV-induced mutagenesis.(8) We assayed mutagenesis in untreated cells and cells treated with a 40 J/m^2^ dose of 254 nm UV-C light, matching prior literature reports, by quantifying the rate of resistance to the drug rifampicin (Rif^R^) (Figure 2A and Table S1). Consistent with these prior reports, the Rif^R^ rate for the wild-type (WT) strain was approximately one in 10^8^ in untreated cells and increased by 15-fold in UV-treated cells (*p* < 0.05). Deletion of either Pol Y1 or Pol Y2 alone led to a 3-fold reduction in UV-mutagenesis relative to WT (*p* < 0.05), broadly consistent with the results of Sung, et al.(7) Notably, deletion of both Pol Y1 and Pol Y2 eliminated any UV-induced mutagenesis (*p* < 0.05), indicating that Pol Y1 and Pol Y2 act in separate pathways to generate mutations upon UV treatment.

**Figure 2.**
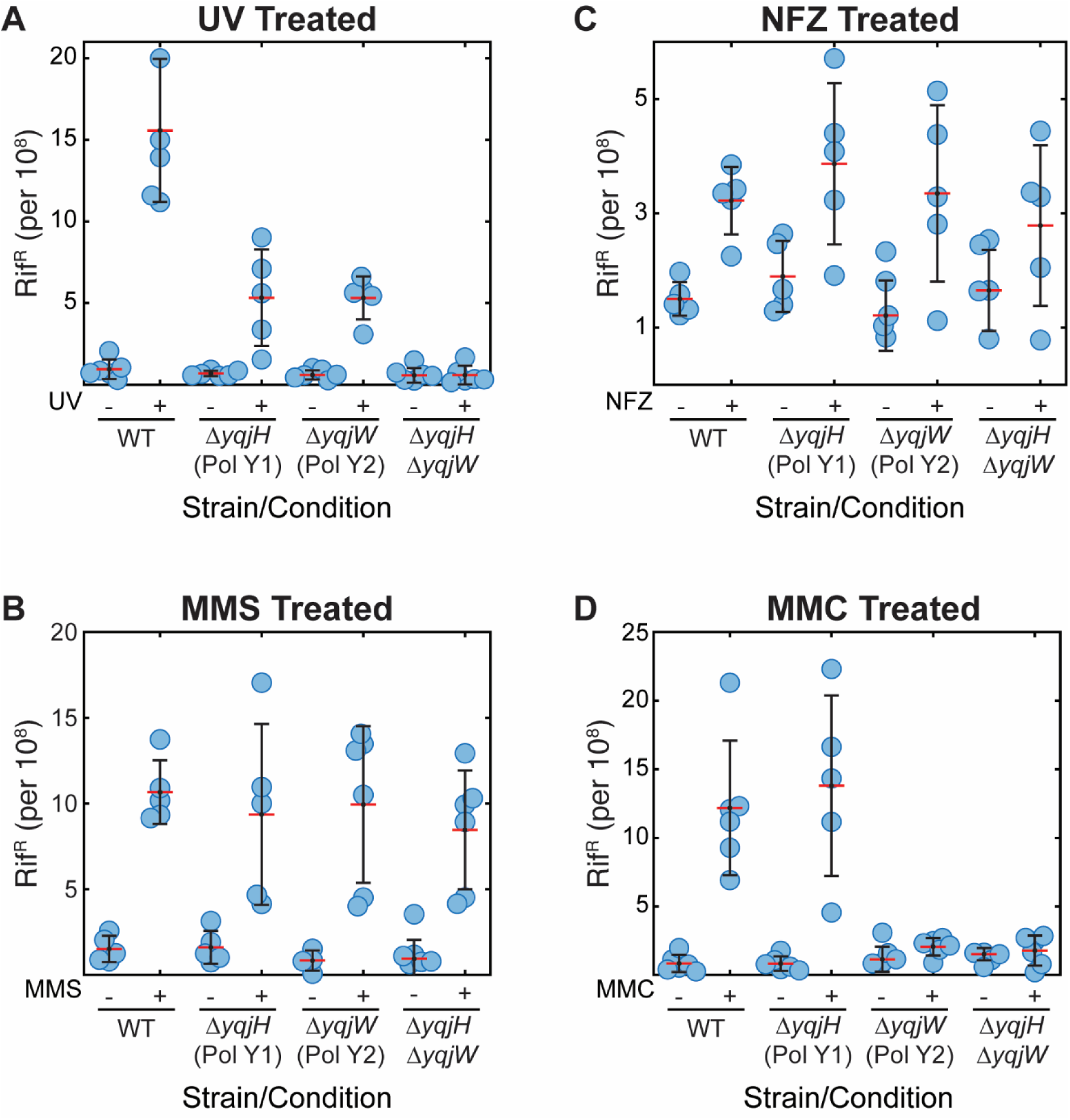
Relative mutagenesis rates for *B. subtilis* strains in the presence of different types of DNA damage. Proportion of rifampicin resistant (Rif^R^) cells for WT, ΔPol Y1 knockout, ΔPol Y2 knockout, and ΔPol Y1 ΔPol Y2 double knockout strains in untreated cells and after treatment with (A) 40 J/m^2^ 254 nm UV light, (B) 10 mM MMS, (C) 100 µM NFZ, and (D) 200 ng/mL MMC. For each dataset, the individual replicates are shown as circles, the red line represents the mean value, and the error bars represent the standard deviation of at least five replicates.

Next, we measured mutagenesis induced by MMS treatment. In *E. coli*, there is a modest reduction in mutagenesis in the absence of Pol V, but a substantial (approximately order of magnitude) increase in mutagenesis upon Pol IV deletion, indicating that Pol V performs highly error-prone synthesis in the absence of Pol IV.(22) We assayed mutagenesis in cells treated with 10 mM MMS for 1 h for the same set of four strains (Figure 2B and Table S1). We found that the Rif^R^ rate for all strains increased by approximately 5 – 10-fold upon MMS treatment, to approximately one in 10^7^ (*p* < 0.05). However, although cells lacking Pol Y1 were sensitized to MMS treatment in survival assays, we found no effect of either Pol Y1 or Pol Y2 deletion on MMS-induced mutagenesis. Likewise, the double Pol Y1 and Pol Y2 knockout had an MMS-induced mutation rate that was statistically indistinguishable from WT. Thus, although Pol Y1 acts to confer tolerance to MMS-induced DNA damage, it does not contribute to mutagenesis.

In *E. coli*, NFZ was reported to induce mutagenesis more weakly than MMS.(31) Surprisingly, although Pol IV promotes survival to NFZ treatment, it has no impact on NFZ-induced mutagenesis.(32) To our knowledge, no prior work has studied the effect of Pol Y1 and Pol Y2 on NFZ-induced mutagenesis in *B. subtilis*. We quantified mutagenesis in cells treated with 100 µM NFZ for 1 h (Figure 2C and Table S1). For the WT strain, there was a small (approximately 2-fold) but statistically significant (*p* < 0.05) increase in the Rif^R^ rate upon NFZ treatment; the same was true for the Pol Y1 knockout strain. Although similar increases were also observed for the Pol Y2 knockout strain and the double knockout, they did not reach statistical significance at the *p* < 0.05 level. However, the NFZ-induced Rif^R^ rate was statistically indistinguishable for all three knockout strains relative to WT, indicating no effect of Pol Y1 or Pol Y2 on NFZ-induced mutagenesis; these results are consistent with the survival assay results showing no effect of Pol Y1 and/or Pol Y2 deletion on tolerance of NFZ-induced DNA damage.

Finally, we assayed mutagenesis upon MMC treatment. As for MMC survival, we are unaware of any studies in *E. coli* on the roles of Pol IV and Pol V in MMC-induced mutagenesis. In *B. subtilis*, both Pol Y1 and Pol Y2 contribute to MMC-induced mutagenesis in sporulating cells,(12) but no studies have investigated their effect in vegetative cells. We assayed mutagenesis in cells treated with 200 ng/mL MMC for 1 h (Figure 2D and Table S1), a condition used previously in other mutagenesis experiments(33) as well as microscopy experiments(34) in *B. subtilis*. For the WT strain, MMC treatment increased the mutation rate by approximately 15-fold (*p* < 0.05); the same was true for the Pol Y1 knockout strain, which was statistically indistinguishable from WT. However, there was a complete loss of MMC-induced mutagenesis for the Pol Y2 deletion, either on its own or in the double knockout strain (*p* > 0.05 for untreated vs. MMC-treated and *p* < 0.05 for MMC-treated relative to WT). Thus, although Pol Y2 does not contribute to survival upon MMC treatment, Pol Y2 is entirely responsible for MMC-induced mutagenesis.

### DNA damage leads to coupled changes in the cellular localization of Pol Y1 and the replisome

Having characterized the effect of Pol Y1 and Pol Y2 deletion on cell survival and mutagenesis upon treatment with different types of DNA damage, we next explored the effect of these treatments on the localization of the replisome and Pol Y1 in live cells (Figure 3, top panel). As a marker for sites of replication, we used the core replisome component DnaX, a subunit of the clamp-loader complex, fused to the YFP variant mYPet.(35) Prior work by us(11,13) and others(36–39) has shown that DnaX forms distinct foci at sites of replication (Figure 3, middle panel). To image single Pol Y1 molecules, we used a C-terminal fusion to the HaloTag, labeled with the Janelia Fluor X 554 (JFX_554_) dye (Figure 3, bottom panel);(40) we validated this imaging strategy previously, showing that the Pol Y1 fusion retains full functionality and that the JFX_554_ labeling procedure has no detectable effects on cell growth or morphology.(11) Consistent with our prior work, we found that untreated cells growing in minimal S7_50_-sorbitol media typically contained one or two DnaX foci (mean ± S.E.M.: 1.76 ± 0.02; Table S2). We calculated the average cellular position of DnaX foci by generating a normalized set of coordinates along the long and short cell axes for each cell (Figure 4A) and found that foci were generally localized at the quarter and three-quarter positions along the long-cell axis (Figure 4B) and at mid-cell along the short-cell axis (Figure S1A), consistent with expectations for normally growing cells. Also consistent with our prior work, we found that Pol Y1 had a similar average localization pattern to DnaX, albeit with a somewhat broader distribution (Figure 4B).(11)

**Figure 3.**
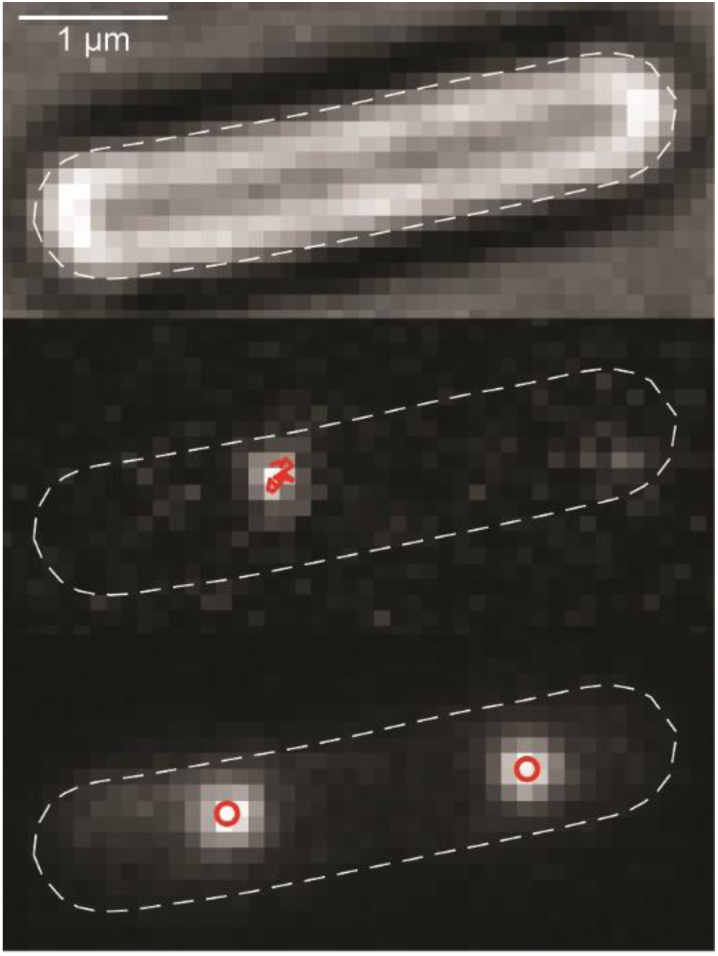
Representative micrographs of Pol Y1 and DnaX. (Top) Transmitted white light micrograph of *B. subtilis* cell with overlaid cell outline and 1 µm scale bar. (Middle) Fluorescence micrograph of single Pol Y1-Halo-JFX_554_ molecule with overlaid trajectory. (Bottom) Fluorescence micrograph of DnaX-mYPet foci with overlaid centroids.

**Figure 4.**
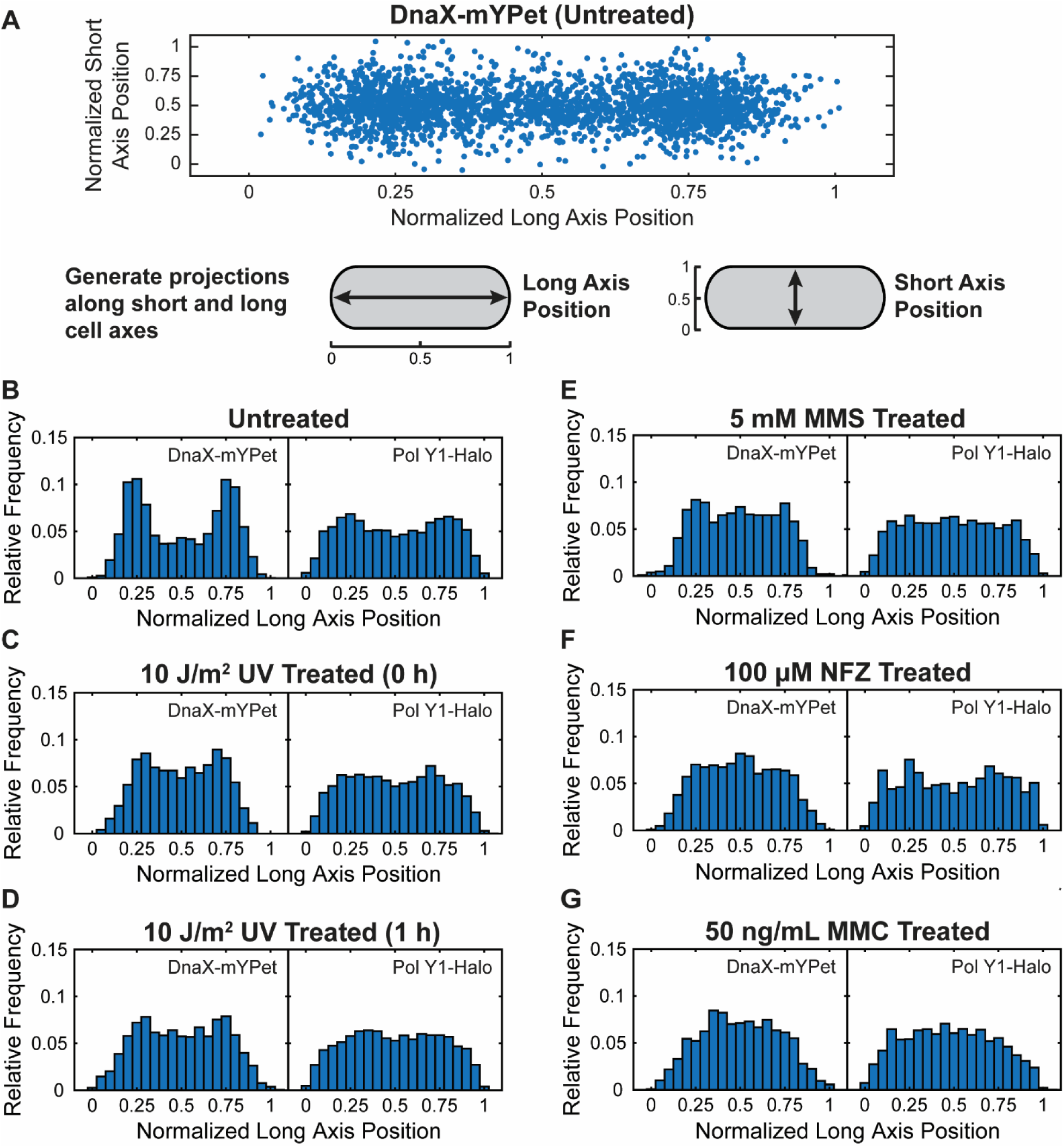
Average cellular localization of DnaX-mYPet and Pol Y1-Halo in the presence of different types of DNA damage. (A) Scatter plot of normalized positions of DnaX foci in untreated cells and cartoon of long and short cell axis projections. Long axis projections of DnaX foci (left) and Pol Y1 trajectories (right) in (B) untreated cells and after treatment with (C) 10 J/m^2^ 254 nm UV light (*t* = 0 h), (D) 10 J/m^2^ 254 nm UV light (*t* = 1 h), (E) 5 mM MMS, (F) 100 µM NFZ, and (G) 50 ng/mL MMC.

Next, we asked how the location of replication sites and Pol Y1 were altered by UV treatment. We tested the effect of treating cells with two different UV doses, 10 and 20 J/m^2^, imaging cells both immediately after exposure and after 1 h of further growth to detect delayed effects of UV treatment. These doses are in a range used in prior imaging studies in *E. coli*.(6,41) However, to determine the effect of these treatments on cell survival in *B. subtilis*, we plated cells before and after UV exposure and quantified the fold-change in the number of CFUs/mL. As reported previously, the number of CFUs/mL doubles (mean ± std.: 2.1 ± 0.4; Table S3) under these growth conditions in the absence of any perturbations. We found that the 10 J/m^2^ UV dose led to a moderate reduction in cell viability immediately after treatment (0.72 ± 0.08), whereas the higher 20 J/m^2^ dose led to a greater reduction (0.4 ± 0.1). However, the number of CFUs/mL approximately doubled after 1 h for both treatments, consistent with the increase observed in untreated cells, indicating a recovery of growth. We also quantified the cell morphology and the average number of DnaX foci per cell after UV treatment, finding that there were moderate increases (approximately 20%) in cell length and in the number of DnaX foci for both doses at the 1 h timepoint (Table S2). For the 10 J/m^2^ dose, there were no qualitative changes in DnaX and Pol Y1 localization at either timepoint, with long-axis localization still peaking at the quarter and three-quarter cell positions (Figure 4C and D). However, for the 20 J/m^2^ dose, both DnaX and Pol Y1 localization shifted to mid-cell along the long cell axis (Figure S2A and B), retaining mid-cell localization along the short-cell axis (Figure S1D and E); this shift is consistent with perturbations to replication and was previously observed in cells treated with 4-NQO.(11)

Next, we tested the effect of treating cells with three different concentrations of MMS (5, 10, and 15 mM) for 1 h in liquid culture; these concentrations are in a range previously used in imaging studies in *E. coli*(42) and *B. subtilis*.(43) The lowest concentration (5 mM) slowed growth but did not lead to a loss of cell viability (1.4 ± 0.3; Table S3), the intermediate concentration (10 mM) led to a modest reduction in viability (0.9 ± 0.1), and the highest concentration (15 mM) led to a substantial reduction (0.30 ± 0.03). There were minimal changes in cell length (< 10% reduction) for all three concentrations (Table S2). Although there was no change in the number of DnaX foci per cell at the lowest concentration, there were moderate (15 – 20%) reductions at the higher two concentrations. All three treatments led to a coupled shift in both DnaX and Pol Y1 localization toward mid-cell along the long-cell axis, but with the shift becoming more pronounced as MMS concentration increased (Figure 4E and Figure S2C and D).

For NFZ, we found that treatment with a wide range of concentrations from 50 to 250 µM slowed growth but did not lead to substantial loss of viability (Table S3); as expected, treatment with the dimethylformamide (DMF) solvent alone had no effect. Thus, we decided to test exposure to two concentrations, 50 and 100 µM, for 1 h in liquid culture, both of which stalled growth without substantial cell killing (1.1 ± 0.3 and 1.0 ± 0.1, respectively); these conditions are similar or identical to NFZ treatments used in prior imaging studies of *E. coli* Pol IV.(4,44) The higher concentration led to a modest (approximately 15% reduction) in cell length and somewhat larger (approximately 15 – 25%) reductions in the number of DnaX foci per cell (Table S2). For the 50 µM NFZ concentration, there was a shift in both DnaX and Pol Y1 localization toward mid-cell along the long cell axis (Figure S2E); for the 100 µM concentration, there was a similar shift for DnaX, although less so for Pol Y1 (Figure 4F).

Finally, we tested the effect of treating cells with three different concentrations of MMC (50, 100, and 200 ng/mL) for 1 h in liquid culture. The lower two concentrations were used in prior imaging studies of *B. subtilis*,(45,46) whereas the higher concentration was used in a prior mutagenesis(33) and imaging experiments.(34) These concentrations lead to increasing loss of viability (0.8 ± 0.2, 0.5 ± 0.1, and 0.18 ± 0.07, respectively; Table S3) (Table S3). We confirmed that treatment with the dimethyl sulfoxide (DMSO) solvent alone did not affect growth. Given the significant reduction in viability at the highest concentration, we decided to use only the lower two concentrations in imaging experiments. Both treatments led to a moderate (∼ 20% increase) in cell length, but no statistically significant changes in the number of DnaX foci per cell (Table S2). The DnaX and Pol Y1 average localization was similar for both treatments (Figure 4G and Figure S2F), with a shift in localization toward mid-cell along the long cell axis, indicative of perturbed replication.

### DNA damage does not induce substantial changes in Pol Y1 colocalization with sites of replication

As described in the previous section, measurements of average cellular localization showed concomitant shifts in the position of replication sites and Pol Y1 upon increasing levels of DNA damage. However, these qualitative comparisons do not allow us to quantify changes in Pol Y1 enrichment at replication sites. Thus, we used the radial distribution function approach to measure the degree of Pol Y1-DnaX colocalization. In brief, the radial distribution function *g*(*r*) reports on the colocalization of two proteins in the cell relative to the colocalization that would be observed by chance due to confinement. The *g*(*r*) value at a given separation distance, or radius, is the fold-enrichment relative to chance of Pol Y1 at that distance from the nearest DnaX focus. Thus, a *g*(*r*) value of 1 indicates no enrichment between the two proteins relative to chance, whereas *g*(*r*) > 1 at short separation distances indicates colocalization.(4,47)

In previous work, we measured the radial distribution function for Pol Y1 and DnaX in untreated cells and found *g*(*r*) ≈ 3 at short distances, indicative of a moderate level of colocalization between Pol Y1 and sites of replication;(11) we repeated these measurements in untreated cells and found *g*(*r*) ≈ 2.0 (Figure 5A and Table S4), in reasonable agreement with our prior report. For *E. coli* Pol IV, diverse types of DNA damage, and even damage-independent replication perturbation by the drug hydroxyurea, were shown to produce large increases, typically around four-fold, in Pol IV colocalization with replication sites, even for non-cognate lesions that Pol IV cannot bypass.(4,6) However, we previously found that there was minimal change in Pol Y1-DnaX colocalization upon treatment with the drug 4-NQO.(11)

**Figure 5.**
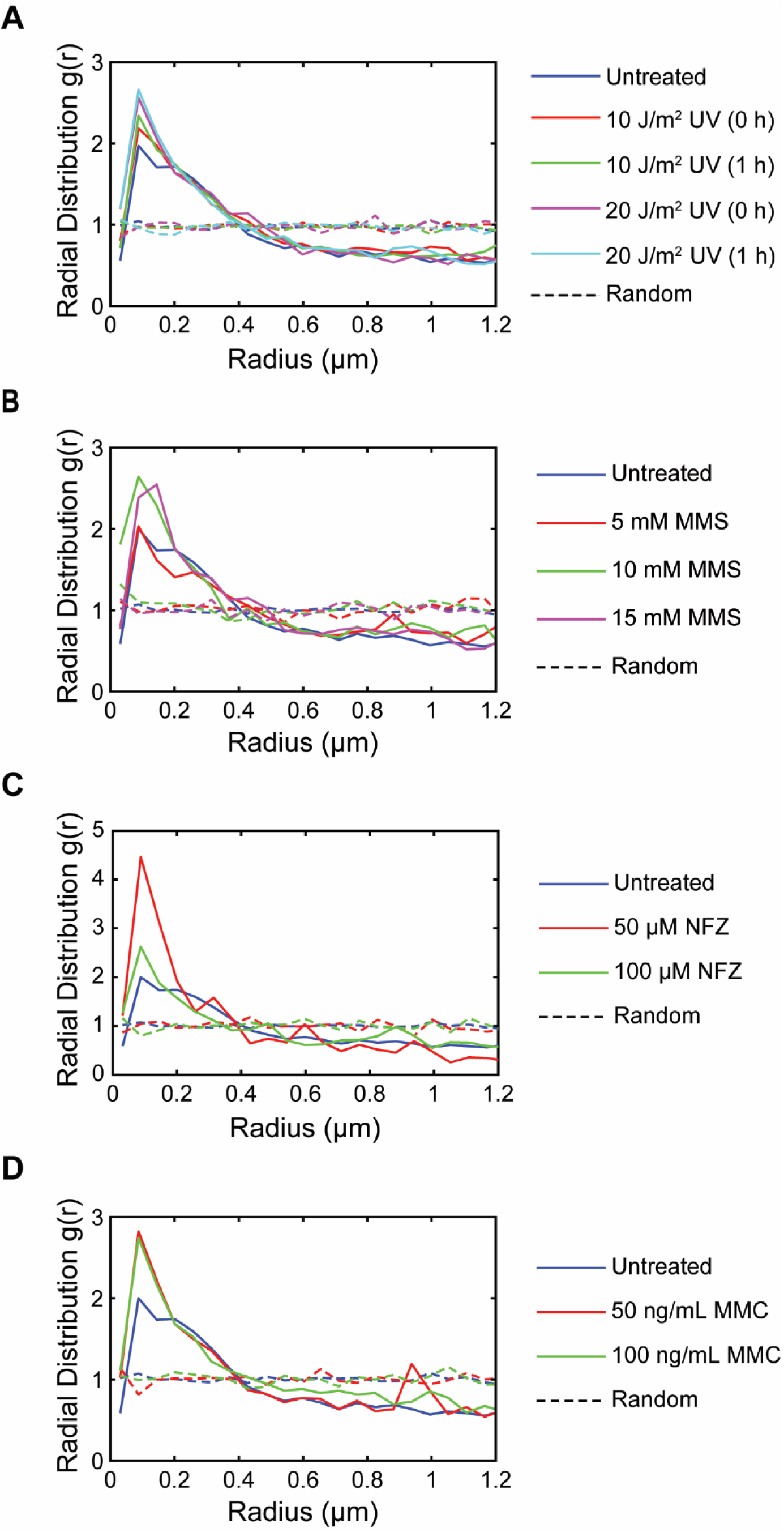
Radial distribution function *g*(*r*) analysis of Pol Y1-Halo and DnaX-mYPet colocalization in the presence of different types of DNA damage. Pol Y1-DnaX *g*(*r*) in untreated cells and (A) after treatment with 10 and 20 J/m^2^ 254 nm UV light (*t* = 0 and 1 h), (B) 5, 10, and 15 mM MMS, (C) 50 and 100 µM NFZ, and (D) 50 and 100 ng/mL MMC. Random *g*(*r*) curves are shown as dashed lines. Values of *g*(*r*) > 1 indicates colocalization, whereas *g*(*r*) = 1 indicates no colocalization. Note different *y*-axis scale in panel (C).

To test whether this lack of damage-induced enrichment is a general phenomenon, we measured the Pol Y1-DnaX radial distribution function for all the different DNA damage treatments described previously. For UV treatment, both 10 and 20 J/m^2^ dosages produced only modest increases in Pol Y1-DnaX colocalization at both timepoints; the increases for 20 J/m^2^ were marginally higher, with a maximum *g*(*r*) ≈ 2.7 (Figure 5A). For MMS treatment, the *g*(*r*) curve for the lowest concentration (5 mM) was very similar to the untreated curve, whereas the 10 and 15 mM concentrations led to a slight increase in colocalization (maximum *g*(*r*) ≈ 2.6) (Figure 5B).

Results for NFZ were more mixed; there was a marked increase in colocalization (maximum *g*(*r*) ≈ 4.6) but only for the lower concentration (50 µM), whereas the *g*(*r*) curve for the higher concentration (100 µM) was very similar to the untreated curve (Figure 5C). Finally, both MMC treatments led to a similar modest increase in Pol Y1-DnaX colocalization (maximum *g*(*r*) ≈ 2.8) (Figure 5D). Taken together, these results indicate that Pol Y1 is not strongly recruited to replication sites upon DNA damage. Further, they reveal no obvious differences in Pol Y1 enrichment between cognate (UV, MMS) and non-cognate (NFZ, MMC) DNA damaging agents.

### DNA damage has differing effects on Pol Y1 mobility in the cell

In previous studies of protein mobility in the cell, DNA polymerases and other DNA-binding proteins have been found to exist in two broad populations: a static population, immobilized via interactions with DNA or with other DNA-bound proteins, and a mobile population, diffusing throughout the cell.(4,48) Although not all static polymerase molecules are actively performing DNA synthesis, it is expected that molecules synthesizing DNA are found in this fraction. Previously, we characterized the static and mobile populations of Pol Y1 and found that static Pol Y1 molecules are enriched near sites of replication in the cell during normal growth.(11) In *E. coli*, DNA damage has been shown to increase the static population of Pol IV(4) and of DNA repair proteins like Pol I and ligase;(48) these static molecules are presumably binding at or near DNA lesions.

Before investigating the effect of DNA damage on Pol Y1 mobility, we first characterized Pol Y1 mobility in untreated cells. Following our previous approach, we calculated the apparent diffusion coefficient (*D*\*) from the mean squared displacement of each trajectory (Figure 6A and Table S5). We then fit the resulting *D*\* distribution to a 3-population model; as we found previously,(11) and as others have seen for other DNA-binding proteins,(49) a 2-population model with a single static and mobile population does not adequately fit the distribution. The resulting fit yields a static population with low mobility (*D*\* ≈ 0.08 µm^2^/s), a highly mobile population (*D*\* ≈ 1.0 µm^2^/s), and an intermediate population (*D*\* ≈ 0.2 µm^2^/s). Consistent with our prior study, we found that static molecules represented ∼ 30% of the population (32 ± 3% vs. 28 ± 3% previously) whereas mobile molecules represented almost half of the population (48 ± 3% vs. 47 ± 3% previously).(11)

**Figure 6.**
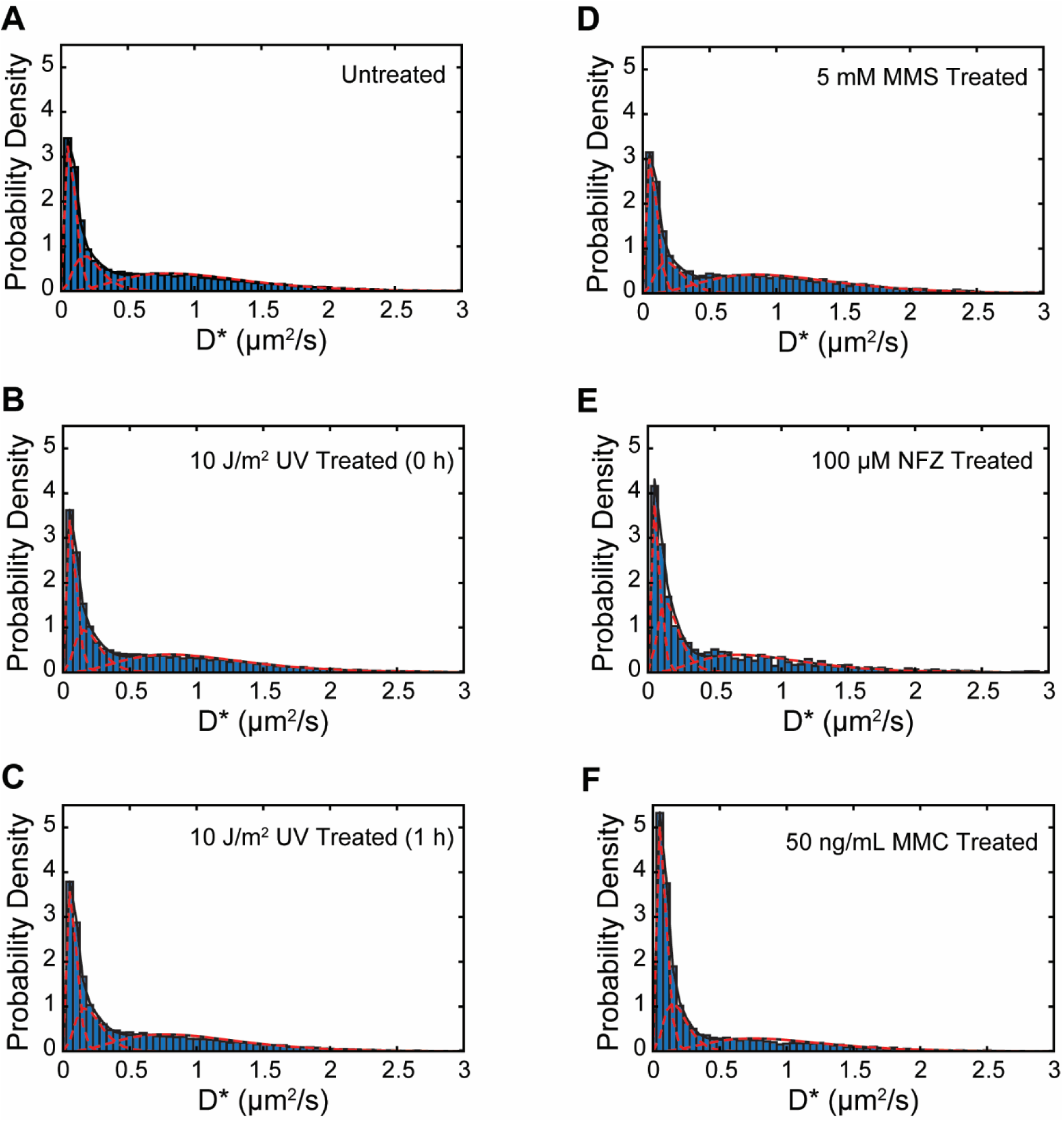
Apparent diffusion coefficient (*D*\*) distributions and corresponding three-population fits for Pol Y1-Halo in the presence of different types of DNA damage. Pol Y1 *D*\* distributions in (A) untreated cells and after treatment with (B) 10 J/m^2^ 254 nm UV light (*t* = 0 h), (C) 10 J/m^2^ 254 nm UV light (*t* = 1 h), (D) 5 mM MMS, (E) 100 µM NFZ, and (F) 50 ng/mL MMC. Individual populations are shown as red dashed lines and overall fits are shown as solid black lines.

Next, we asked whether treatment with DNA damaging agents altered the mobility of Pol Y1 molecules in the cell. For UV treatment, the 10 J/m^2^ dosage produced only minimal changes in Pol Y1 diffusion at both timepoints (Figure 6B and C). The 20 J/m^2^ dosage produced a modest (∼ 5 – 10%) increase in the static population with a corresponding decrease in the mobile population at the *t* = 0 timepoint, with a larger (∼ 20%) decrease in the mobile population at the *t* = 1 h timepoint (Figure S3A and B). MMS treatment had only minor effects on Pol Y1 mobility. Both the lowest (5 mM) and highest (15 mM) concentrations led to a modest (∼ 10%) decrease in the static population, whereas the intermediate concentration (10 mM) had no effect (Figure 6D and Figure S3C and D,). In contrast, NFZ treatment led to larger changes in Pol Y1 mobility than either UV or MMS. For the lower concentration (50 µM), there was little change in the static population but a larger increase in the intermediate population (∼ 70%), and a decrease in the mobile population (∼ 30%) (Figure S3E). For the higher concentration (100 µM), the direction of change was the same for all populations, but the magnitudes were generally smaller (Figure 6E).

Like NFZ treatment, MMC led to larger changes in Pol Y1 diffusion, but without an obvious dose-dependent effect. The lower concentration (50 ng/mL) produced a larger increase in the static population (∼ 40%), paired with a corresponding decrease in the mobile population (∼ 30%) (Figure 6F). The higher concentration (100 ng/mL) led to a similar decrease in the mobile population but with a smaller (∼ 10%) increase in the static population; a larger increase (∼ 50%) in the intermediate population accounted for the remaining share of molecules (Figure S3F). Thus, although changes in diffusion were larger than changes in Pol Y1 enrichment at replication forks, the results were not consistent across different types of DNA damage, and even the largest effects were not substantial.

## Discussion

*E. coli* has served as the model system for bacterial TLS since the identification of translesion polymerases a quarter century ago,(1,2) yet it remains poorly understood how widely the mechanisms elucidated in *E. coli* hold in other species. In a previous study, we investigated the subcellular localization and dynamics of the *B. subtilis* TLS polymerase Pol Y1, the homolog of *E. coli* Pol IV, both during normal growth and after treatment with the cognate DNA damaging agent 4-NQO.(11) We also characterized the effect of deleting Pol Y1 or Pol Y2, the *E. coli* Pol V homolog, on 4-NQO survival and damage-induced mutagenesis, finding that Pol Y1 contributed to survival whereas neither polymerase had a significant effect on mutagenesis. In this study, we have expanded our investigation to include a range of different DNA damaging agents (UV, MMS, NFZ, and MMC) with the goal of broadening our understanding of TLS mechanisms in *B. subtilis*.

Because there is little or no literature data on treatment with these drugs in *B. subtilis*, we first determined the effect of Pol Y1 and Pol Y2 deletion, individually or in combination, on cell survival after treatment with different doses of these four damaging agents. These experiments revealed several differences in comparison to *E. coli*. Consistent with prior reports,(7,10,13) we found that Pol Y1 promoted damage survival upon UV treatment, with an approximately 10-fold reduction in survival upon Pol Y1 deletion, whereas deletion of Pol Y2 had no effect; these results are contrary to *E. coli*, where Pol V alone promotes survival to UV lesions.(14,15) They are also in contrast to a prior study with sporulating *B. subtilis* cells, in which both Pol Y1 and Pol Y2 contributed to survival.(12) We found that Pol Y1 promoted survival to MMS, although Pol Y1 deletion had a relatively modest effect (2 – 3-fold reduction in survival relative to WT); these findings are broadly consistent with *E. coli*, where Pol IV promotes MMS survival, although ΔPol IV cells are more highly sensitized to MMS treatment.(22) Results for NFZ treatment provided another point of difference with *E. coli*. We found that neither Pol Y1 nor Pol Y2 contributed to NFZ tolerance; in contrast, NFZ lesions are well-characterized as cognate lesions for Pol IV.(15) These results illustrate differences in the roles of Pol Y1 and Pol Y2 and their *E. coli* homologs; given the modest effect of Pol Y1 deletion on MMS survival and the lack of effect observed for NFZ and MMC, they may also suggest that other pathways in *B. subtilis* play a greater role in DNA damage tolerance than TLS does. Notably, we found no effect of either Pol Y1 or Pol Y2 deletion on MMC survival, in contrast to a prior report for sporulating cells, in which both contributed to survival,(12) suggesting that TLS mechanisms may differ during vegetative growth and sporulation in *B. subtilis*.

DNA damage induces mutagenesis by upregulation of error-prone pathways like TLS; in *E. coli*, Pol IV and Pol V have both been found to contribute to damage-induced mutagenesis under different conditions. Thus, in addition to characterizing the roles of Pol Y1 and Pol Y2 in the tolerance of different types of DNA damage, we also explored their effects on mutagenesis using a rifampicin resistance assay. Consistent with prior studies, we found that deletion of Pol Y1 or Pol Y2 had no measurable effect on the spontaneous mutagenesis rate.(7,11) For a UV treatment condition that increased mutagenesis by approximately 15-fold for WT, we found that deletion of Pol Y1 and Pol Y2 individually led to a 3-fold reduction in mutagenesis, whereas deletion of both eliminated UV-induced mutagenesis entirely. However, for an MMS treatment condition that increased mutagenesis by approximately 10-fold for WT, we found no effect of Pol Y1 and/or Pol Y2 deletion on mutagenesis, in contrast to the modest effect on cell survival observed for a Pol Y1 deletion. For NFZ, we found that treatment only increased the mutagenesis rate by 2-fold, consistent with the relatively modest impact that NFZ has on mutagenesis in *E. coli*.(32) Under these conditions, we observed no effect of Pol Y1 and/or Pol Y2 deletion on mutagenesis, mirroring the lack of impact of these deletions on NFZ survival. Finally, for MMC, we tested a treatment condition that increased mutagenesis by approximately 15-fold for WT; we found that Pol Y1 deletion had no impact on mutagenesis, but Pol Y2 deletion, either alone or in combination with Pol Y1 deletion, eliminated MMC-induced mutagenesis. These results are surprising given that Pol Y2 deletion had no effect on cell survival; however, a similar phenomenon has been observed for MMS treatment in *E. coli*, for which Pol V deletion has no impact on survival but does reduce mutagenesis.(22) Taken together, these data reveal further differences in the roles of Pol Y1 and Pol Y2 and their E*. coli* homologs in damage-induced mutagenesis. They also indicate that TLS polymerases can make distinct contributions to survival and mutagenesis for the same type of DNA damage.

In *E. coli*, single-molecule imaging studies have suggested a spatial component of Pol IV regulation; in brief, Pol IV is not highly enriched near sites of replication during normal growth, but it is strongly recruited upon DNA damage or other replication perturbations, even for non-cognate lesions or in other situations where Pol IV cannot resolve the challenge to replication.(4,6) In our prior study, we found that Pol Y1 was moderately enriched near replication sites even during normal growth, in contrast to Pol IV; we also found that Pol Y1 was not further enriched upon 4-NQO treatment, again in contrast to Pol IV.(11) Likewise, 4-NQO treatment had no impact on Pol Y1 mobility. In this study, we asked whether this lack of damage-induced enrichment was unique to 4-NQO treatment or consistent for other types of DNA damage. We imaged cells after treatment with different doses of UV, MMS, NFZ, and MMC, choosing treatment conditions that halted cell growth or produced a moderate reduction in cell viability.

In general, we saw only minimal changes in the degree of Pol Y1 enrichment at replication sites upon different types of DNA damage. The only exception was NFZ treatment, for which we observed a moderate increase in colocalization for the 50 µM concentration. Interestingly, of the different damaging agents tested in similar experiments with *E. coli* Pol IV, NFZ was the only drug that did not produce a substantial increase in Pol IV colocalization with sites of replication; this result was interpreted as suggesting that Pol IV performs TLS past NFZ-induced lesions rapidly at the replication fork, without substantial enrichment being required to rescue replication.(4) This model might be consistent with our findings that Pol Y1 is not able to promote survival to NFZ-induced DNA damage, although it is hard to explain why comparable enrichment was not observed at the higher NFZ concentration. Taken together, the results for the four damaging agents are broadly consistent with what we observed previously for 4-NQO treatment and in contrast to the behavior of *E. coli* Pol IV.

Like other DNA-binding proteins, Pol Y1 exists in static and mobile population in the cell, with the former immobilized via interactions with DNA or with other DNA-bound proteins.(4,48) In *E. coli*, DNA damage was found to increase the static fraction of Pol IV molecules by over 2-fold, consistent with Pol IV binding near sites of stalled replication and performing TLS;(4) similar behavior has been observed for DNA repair proteins like the polymerase Pol I and ligase, with approximately 5-fold increases in the static fraction.(48) However, we previously found no effect of 4-NQO treatment on Pol Y1 diffusion.(11) For the damaging agents tested in this study, DNA damage led to more pronounced changes in Pol Y1 diffusion than in its enrichment at replication forks, although the changes depended strongly on the type of DNA damage. UV, MMS, and NFZ led to modest (typically ∼ 10%) or less changes in the static population, with UV and NFZ treatment increasing the static fraction and MMS treatment actually decreasing the static fraction. MMC treatment led to a larger increase in the static fraction (∼ 40%), albeit only for the lower dose (50 ng/mL). However, although this change was larger than those seen for other damaging agents, it is much smaller than the reported fold-changes for Pol IV(4) and Pol I(48) in *E. coli*. Taken together, our results suggest that DNA damage generally increases Pol Y1 binding, although the effects are relatively modest.

This study provides new insight into the activity of TLS polymerases in *B. subtilis*, but it also raises new questions. First, what explains the discrepancies between the contributions of Pol Y1 and Pol Y2 to DNA damage survival and mutagenesis? For example, how does Pol Y1 promote survival to MMS without contributing to mutagenesis, and how does Pol Y2 promote MMC-induced mutagenesis while having no impact on survival? Second, are TLS pathways different in vegetative and sporulating cells? A prior study in sporulating cells found effects of Pol Y1 and Pol Y2 deletion on DNA damage survival and mutagenesis after UV and MMC treatment that differ from our results.(12) Related to the molecular mechanisms of TLS, what explains the lack of damage-induced enrichment of Pol Y1 at replication forks and how is it acting to promote cell survival after damage? Both Pol IV(4,50) and Pol Y1(11) must bind the clamp to perform TLS, and clamp-binding mutations phenocopy a knockout. However, Pol IV is enriched at replication sites upon DNA damage primarily through interactions with the replisome component single-stranded DNA-binding protein (SSB); weakening the Pol IV-SSB interaction leads to a moderate defect in DNA damage tolerance, but a substantial loss in enrichment at the fork.(5,6) It is unknown whether Pol Y1 and Pol Y2 bind SSB; they have not been identified as SSB-interacting proteins via screening,(51) although to our knowledge no study has searched for this interaction specifically. Finally, although Pol Y2 is only present in the cell after SOS induction,(8) does it have similar localization and dynamics to Pol Y1, or is it more strongly recruited to stalled replication forks? Given the apparent diversity of bacterial TLS mechanisms, future work is needed to address these questions and to develop a more comprehensive picture of bacterial TLS.

## Materials and Methods

*Bacterial strain construction:* All bacterial strains used in this study were constructed in the *B. subtilis* WT background PY79.(52,53) Genetic modifications were introduced by transformation of double-stranded DNA fragments generated by PCR and Gibson assembly or by transformation of genomic DNA, followed by selection on appropriate antibiotic plates. Detailed strain information is provided in the Supplementary Methods, including lists of all oligonucleotides (Table S6) and strains (Table S7) used in this study.

*DNA damage survival assays:* Quantitative survival rates were determined for strains treated with different types and doses of DNA damaging agents following previously reported procedures,(11,13) with slight modifications for different damaging agents. In all cases, strains were first streaked out from glycerol stocks onto LB Lennox agar plates containing an antibiotic for selection. The following day, overnight cultures were inoculated from single colonies and incubated overnight in LB Lennox media shaking at 22 °C. The next morning, fresh cultures were inoculated and grown to exponential phase shaking at 37 °C, then serially diluted and plated on LB Lennox agar plates as described below. After overnight incubation at 37 °C, colonies were enumerated, and the survival rate was calculated. For NFZ treatment, colonies were small and difficult to count accurately after overnight incubation at 37 °C. Thus, plates were left at room temperature for an additional 24 h before counting.

Survival to UV light was assayed exactly as described previously(13) by plating cells on plain LB Lennox agar plates and irradiating with varying doses of 254 nm UV-C light (Analytik Jena UVS-28 EL). Survival to MMS, NFZ, and MMC ware assayed following the same procedure described previously for 4-NQO treatment.(11) In brief, LB Lennox agar plates were prepared containing different concentrations of drug, with the total volume of solvent added held constant across all drug concentrations. MMS was used as a neat liquid, NFZ solutions were prepared in DMF, and MMC solutions were prepared in DMSO.

At least three independent replicates were performed for each damaging agent and checked for consistency. For MMS, we noted higher than normal variability in the survival rates. Therefore, we conducted a total of nine replicates. Inconsistent data points were identified using the MATLAB function isoutlier using the default parameters and were removed from the dataset; in all cases, at least six replicates remained. Although outlier removal reduced the resulting standard deviation in the dataset, it did not have an impact on the qualitative results.

*Rifampicin resistance mutagenesis assays:* Mutagenesis was quantified under different treatment conditions by measuring the rate of resistance to the antibiotic rifampicin (Rif^R^). These assays were performed following previously reported procedures,(11,13) with slight modifications for different damaging agents. UV mutagenesis assays were performed exactly as described previously, by resuspending cells in transparent MgSO_4_ solution and irradiating with a dose of 40 J/m^2^, then resuspending in fresh LB Lennox media and growing overnight.(13) Mutagenesis assays with MMS, NFZ, and MMC followed the same procedure described previously for 4-NQO treatment(11) but adapted for the different drugs. In brief, each drug was added to liquid culture at final concentrations of 10 mM for MMS (from the neat liquid), 100 µM for NFZ (from a 100 mM stock), and 200 ng/mL for MMC (from a 100 µg/mL stock). Cultures were incubated shaking at 37 °C for 1 h and then washed, resuspended in fresh media, and grown overnight exactly as described previously for 4-NQO mutagenesis assays. For all assays, the overnight cultures were serially diluted and plated on LB Lennox agar plates with or without Rif. After overnight incubation, colonies were enumerated, the number of CFUs/mL was determined for Rif^+^ and Rif^−^ plates, and the Rif^R^ rate was calculated.

At least five independent replicates were performed for each damaging agent and checked for consistency. For each damaging agent, an independent dataset was collected in parallel with untreated cells to measure the rate of spontaneous rifampicin resistance Rif^R^.

*Cell culture and sample preparation for microscopy:* Cell culture and sample preparation for microscopy was performed exactly as described previously.(11) Imaging cultures were grown in minimal S7_50_-sorbitol media at 25 mL scale shaking at 37 °C. Samples were harvested in early exponential phase (OD_600nm_ ≈ 0.15 for untreated samples), labeled with JFX_554_ dye at a final concentration of 2.5 nM, and incubated shaking at 37 °C for 15 min. After labeling, cells were washed, concentrated by centrifugation, deposited on an agarose pad containing growth media, and sandwiched between two cleaned coverslips.

*Sample treatment for microscopy:* Cultures were treated with DNA damaging agents when they reached the same OD_600nm_ ≈ 0.15. For UV treatment, the entire culture was transferred to a sterile Petri dish and irradiated with 254 nm UV-C light (Analytik Jena UVS-28 EL) with gentle stirring. An aliquot was taken for labeling and imaging (the *t* = 0 h sample) and the remaining culture was returned to a culture tube and grown for an additional hour shaking at 37 °C, at which point a second aliquot was taken for labeling and imaging (the *t* = 1 h sample). Treatment with MMS, NFZ, and MMC was performed following the same procedure described previously for 4-NQO treatment.(11) In brief, the drug was added as a neat liquid (for MMS) or from a stock solution at a 1:1,000 dilution (for NFZ and MMC) to the culture, after which the culture was grown for an additional hour before a sample was harvested for labeling and imaging.

For all treatment conditions, the effect on cell survival was quantified as described previously for 4-NQO treatment.(11) Aliquots were taken before and after treatment, serially diluted, and plated on plain LB Lennox agar plates. The number of CFUs/mL was determined for treated and untreated samples after overnight incubation and the fold-changed was calculated.

*Microscopy:* Microscopy was performed using a custom fluorescence microscope described previously.(11) A Nikon Ti2-E inverted microscope was equipped with a Nikon CFI Apo 100×/1.49 NA total internal reflection fluorescence (TIRF) objective lens and Chroma filters. Laser excitation (Coherent Sapphire) at 514 nm (∼1 W cm^−2^ power density) and 561 nm (∼15 W cm^−2^ power density) was used to excite DnaX-mYPet and Pol Y1-Halo-JFX_554_ fluorescence, respectively. All movies were recorded using a short integration time (13.9 ms) with a Hamamatsu ImageEM C9100-23BKIT EMCCD camera. Highly inclined thin illumination, or near-TIRF, illumination was achieved by focusing the laser light to the back focal plane of the objective.(54) Brightfield images of cells were recorded for each field of view using white light transillumination.

*Image and data analysis:* Quantitative image analysis was performed exactly as described previously.(11) In brief, the MATLAB package MicrobeTracker(55) was used for cell segmentation of brightfield images, and the MATLAB package u-track(56,57) was used for spot detection and tracking of fluorescence movies. For analysis of DnaX-mYPet foci, the first 20 frames of 514 nm excitation were averaged.

Likewise, the same analysis approaches and custom code were used for determination of average DnaX and Pol Y1 cellular localization, Pol Y1-DnaX colocalization (through radial distribution function analysis), and Pol Y1 diffusion (using the MSD approach to determine the apparent diffusion coefficient *D*\*) as described previously.(11) Cell morphology was determined from MicrobeTracker cell segmentation output.

All imaging experiments were performed on at least two separate days with at least three separate replicates, defined as separate imaging cultures. The exact number of imaging days, replicates, cells, and tracks or foci for all figures are listed in Table S8.

For mutagenesis and imaging datasets, statistical comparisons were made using the two-tailed Wilcoxon rank sum test, implemented in the MATLAB function ranksum.

*Data availability:* Data and custom MATLAB analysis code from this study have been deposited in a Zenodo repository (DOI: 10.5281/zenodo.19355396).

## Acknowledgments

We acknowledge Madeline Drucker, Carolyn Greenwald, and Sarah Rancic for assistance with strain construction. This work was supported by the National Institute of General Medical Sciences of the National Institutes of Health [award number R15GM151677 to E.S.T.] Additional support was provided by the Fordham College at Rose Hill Undergraduate Research Grant program [awards to S.R.M., A.P.C., and I.E.Z.J.], the Fordham University Clare Boothe Luce program [awards to C.M.S. and M.E.M.], and the Len Blavatnik STEM Fellowship [award to M.E.M.]. The funders had no role in study design, data collection and analysis, decision to publish, or preparation of the manuscript.

## References

1. Fuchs RP, Fujii S. Translesion DNA Synthesis and Mutagenesis in Prokaryotes. Cold Spring Harbor Perspectives in Biology. 2013 Dec 1;5(12):a012682–a012682. doi:10.1101/cshperspect.a012682

2. Fujii S, Fuchs RP. A Comprehensive View of Translesion Synthesis in Escherichia coli. Microbiol Mol Biol Rev. 2020 Aug 19;84(3):e00002–20. doi:10.1128/MMBR.00002-20

3. Joseph AM, Badrinarayanan A. Visualizing mutagenic repair: novel insights into bacterial translesion synthesis. FEMS Microbiology Reviews. 2020 Sep 1;44(5):572–82. doi:10.1093/femsre/fuaa023

4. Thrall ES, Kath JE, Chang S, Loparo JJ. Single-molecule imaging reveals multiple pathways for the recruitment of translesion polymerases after DNA damage. Nat Commun. 2017 Dec 18;8(1):2170. doi:10.1038/s41467-017-02333-2

5. Chang S, Thrall ES, Laureti L, Piatt SC, Pagès V, Loparo JJ. Compartmentalization of the replication fork by single-stranded DNA-binding protein regulates translesion synthesis. Nat Struct Mol Biol. 2022 Sep;29(9):932–41. doi:10.1038/s41594-022-00827-2

6. Thrall ES, Piatt SC, Chang S, Loparo JJ. Replication stalling activates SSB for recruitment of DNA damage tolerance factors. Proc Natl Acad Sci USA. 2022 Oct 11;119(41):e2208875119. doi:10.1073/pnas.2208875119

7. Sung HM, Yeamans G, Ross CA, Yasbin RE. Roles of YqjH and YqjW, homologs of the *Escherichia coli* UmuC/DinB or Y superfamily of DNA polymerases, in stationary-phase mutagenesis and UV-induced mutagenesis of *Bacillus subtilis*. J Bacteriol. 2003 Apr;185(7):2153–60. doi:10.1128/JB.185.7.2153-2160.2003 PubMed PMID: 12644484; PubMed Central PMCID: PMC151490.

8. Duigou S, Ehrlich SD, Noirot P, Noirot-Gros MF. Distinctive genetic features exhibited by the Y-family DNA polymerases in *Bacillus subtilis*. Mol Microbiol. 2004 Oct;54(2):439–51. doi:10.1111/j.1365-2958.2004.04259.x PubMed PMID: 15469515.

9. Lenhart JS, Schroeder JW, Walsh BW, Simmons LA. DNA Repair and Genome Maintenance in Bacillus subtilis. Microbiol Mol Biol Rev. 2012 Sep;76(3):530–64. doi:10.1128/MMBR.05020-11

10. Million-Weaver S, Samadpour AN, Moreno-Habel DA, Nugent P, Brittnacher MJ, Weiss E, et al. An underlying mechanism for the increased mutagenesis of lagging-strand genes in Bacillus subtilis. Proc Natl Acad Sci U S A. 2015 Mar 10;112(10):E1096–1105. doi:10.1073/pnas.1416651112 PubMed PMID: 25713353; PubMed Central PMCID: PMC4364195.

11. Marrin ME, Foster MR, Santana CM, Choi Y, Jassal AS, Rancic SJ, et al. The translesion polymerase Pol Y1 is a constitutive component of the *B. subtilis* replication machinery. Nucleic Acids Research. 2024 Sep 9;52(16):9613–29. doi:10.1093/nar/gkae637

12. Rivas-Castillo AM, Yasbin RE, Robleto E, Nicholson WL, Pedraza-Reyes M. Role of the Y-Family DNA Polymerases YqjH and YqjW in Protecting Sporulating Bacillus subtilis Cells from DNA Damage. Curr Microbiol. 2010 Apr;60(4):263–7. doi:10.1007/s00284-009-9535-3

13. O’Neal LG, Drucker MN, Lai NK, Clemente AF, Campbell AP, Way LE, et al. The *B. subtilis* replicative polymerases bind the sliding clamp with different strengths to tune their activity in DNA replication. Nucleic Acids Research. 2025 Jul 19;53(14):gkaf721. doi:10.1093/nar/gkaf721

14. Courcelle CT, Belle JJ, Courcelle J. Nucleotide Excision Repair or Polymerase V-Mediated Lesion Bypass Can Act To Restore UV-Arrested Replication Forks in *Escherichia coli*. J Bacteriol. 2005 Oct 15;187(20):6953–61. doi:10.1128/JB.187.20.6953-6961.2005

15. Ona KR, Courcelle CT, Courcelle J. Nucleotide Excision Repair Is a Predominant Mechanism for Processing Nitrofurazone-Induced DNA Damage in *Escherichia coli*. J Bacteriol. 2009 Aug;191(15):4959–65. doi:10.1128/JB.00495-09

16. Santos-Escobar F, Leyva-Sánchez HC, Ramírez-Ramírez N, Obregón-Herrera A, Pedraza-Reyes M. Roles of *Bacillus subtilis* RecA, Nucleotide Excision Repair, and Translesion Synthesis Polymerases in Counteracting Cr(VI)-Promoted DNA Damage. Henkin TM, editor. J Bacteriol. 2019 Apr 15;201(8). doi:10.1128/JB.00073-19

17. Au N, Kuester-Schoeck E, Mandava V, Bothwell LE, Canny SP, Chachu K, et al. Genetic Composition of the *Bacillus subtilis* SOS System. J Bacteriol. 2005 Nov 15;187(22):7655–66. doi:10.1128/JB.187.22.7655-7666.2005

18. Sinha RP, Häder DP. UV-induced DNA damage and repair: a review. Photochem Photobiol Sci. 2002 Apr;1(4):225–36. doi:10.1039/b201230h

19. Tang M, Pham P, Shen X, Taylor JS, O’Donnell M, Woodgate R, et al. Roles of E. coli DNA polymerases IV and V in lesion-targeted and untargeted SOS mutagenesis. Nature. 2000 Apr;404(6781):1014–8. doi:10.1038/35010020

20. Wyatt MD, Pittman DL. Methylating Agents and DNA Repair Responses: Methylated Bases and Sources of Strand Breaks. Chem Res Toxicol. 2006 Dec 1;19(12):1580–94. doi:10.1021/tx060164e

21. Sikora A, Mielecki D, Chojnacka A, Nieminuszczy J, Wrzesinski M, Grzesiuk E. Lethal and mutagenic properties of MMS-generated DNA lesions in Escherichia coli cells deficient in BER and AlkB-directed DNA repair. Mutagenesis. 2010 Mar 1;25(2):139–47. doi:10.1093/mutage/gep052

22. Scotland MK, Heltzel JMH, Kath JE, Choi JS, Berdis AJ, Loparo JJ, et al. A Genetic Selection for dinB Mutants Reveals an Interaction between DNA Polymerase IV and the Replicative Polymerase That Is Required for Translesion Synthesis. Matic I, editor. PLoS Genet. 2015 Sep 9;11(9):e1005507. doi:10.1371/journal.pgen.1005507

23. McCalla DR, Reuvers A, Kaiser C. Mode of Action of Nitrofurazone. J Bacteriol. 1970 Dec;104(3):1126–34. doi:10.1128/jb.104.3.1126-1134.1970

24. Tu Y, McCalla DR. Effect of activated nitrofurans on DNA. Biochim Biophys Acta. 1975 Aug 21;402(2):142–9. doi:10.1016/0005-2787(75)90032-5 PubMed PMID: 1100114.

25. Bjedov I, Dasgupta CN, Slade D, Le Blastier S, Selva M, Matic I. Involvement of *Escherichia coli* DNA Polymerase IV in Tolerance of Cytotoxic Alkylating DNA Lesions *in Vivo*. Genetics. 2007 Jul 1;176(3):1431–40. doi:10.1534/genetics.107.072405

26. Iyer VN, Szybalski W. A Molecular Mechanism of Mitomycin Action: Linking of Complementary DNA Strands. Proc Natl Acad Sci USA. 1963 Aug;50(2):355–62. doi:10.1073/pnas.50.2.355

27. Tomasz M. The Mitomycins: Natural Cross-linkers of DNA. In: Neidle S, Waring M, editors. Molecular Aspects of Anticancer Drug-DNA Interactions [Internet]. London: Macmillan Education UK; 1994 [cited 2025 Aug 6]. p. 312–49. Available from: http://link.springer.com/10.1007/978-1-349-13330-7_8 doi:10.1007/978-1-349-13330-7_8

28. Tomasz M, Palom Y. The mitomycin bioreductive antitumor agents: Cross-linking and alkylation of DNA as the molecular basis of their activity. Pharmacology & Therapeutics. 1997 Oct;76(1–3):73–87. doi:10.1016/S0163-7258(97)00088-0

29. Dronkert MLG, Kanaar R. Repair of DNA interstrand cross-links. Mutation Research/DNA Repair. 2001 Sep;486(4):217–47. doi:10.1016/S0921-8777(01)00092-1

30. Gawel D, Maliszewska-Tkaczyk M, Jonczyk P, Schaaper RM, Fijalkowska IJ. Lack of Strand Bias in UV-Induced Mutagenesis in *Escherichia coli*. J Bacteriol. 2002 Aug 15;184(16):4449–54. doi:10.1128/JB.184.16.4449-4454.2002

31. Benson RW, Norton MD, Lin I, Du Comb WS, Godoy VG. An Active Site Aromatic Triad in Escherichia coli DNA Pol IV Coordinates Cell Survival and Mutagenesis in Different DNA Damaging Agents. Marinus MG, editor. PLoS ONE. 2011 May 17;6(5):e19944. doi:10.1371/journal.pone.0019944

32. Jarosz DF, Godoy VG, Delaney JC, Essigmann JM, Walker GC. A single amino acid governs enhanced activity of DinB DNA polymerases on damaged templates. Nature. 2006 Jan;439(7073):225–8. doi:10.1038/nature04318

33. Dupes NM, Walsh BW, Klocko AD, Lenhart JS, Peterson HL, Gessert DA, et al. Mutations in the *Bacillus subtilis* β Clamp That Separate Its Roles in DNA Replication from Mismatch Repair. J Bacteriol. 2010 Jul;192(13):3452–63. doi:10.1128/JB.01435-09

34. Li Y, Chen Z, Matthews LA, Simmons LA, Biteen JS. Dynamic Exchange of Two Essential DNA Polymerases during Replication and after Fork Arrest. Biophysical Journal. 2019 Feb;116(4):684–93. doi:10.1016/j.bpj.2019.01.008

35. Nguyen AW, Daugherty PS. Evolutionary optimization of fluorescent proteins for intracellular FRET. Nat Biotechnol. 2005 Mar;23(3):355–60. doi:10.1038/nbt1066

36. Wang X, Montero Llopis P, Rudner DZ. *Bacillus subtilis* chromosome organization oscillates between two distinct patterns. Proc Natl Acad Sci U S A. 2014 Sep 2;111(35):12877–82. doi:10.1073/pnas.1407461111 PubMed PMID: 25071173; PubMed Central PMCID: PMC4156703.

37. Liao Y, Schroeder JW, Gao B, Simmons LA, Biteen JS. Single-molecule motions and interactions in live cells reveal target search dynamics in mismatch repair. Proc Natl Acad Sci U S A. 2015 Dec 15;112(50):E6898-6906. doi:10.1073/pnas.1507386112 PubMed PMID: 26575623; PubMed Central PMCID: PMC4687589.

38. Mangiameli SM, Veit BT, Merrikh H, Wiggins PA. The Replisomes Remain Spatially Proximal throughout the Cell Cycle in Bacteria. Viollier PH, editor. PLoS Genet. 2017 Jan 23;13(1):e1006582. doi:10.1371/journal.pgen.1006582

39. Hernández-Tamayo R, Oviedo-Bocanegra LM, Fritz G, Graumann PL. Symmetric activity of DNA polymerases at and recruitment of exonuclease ExoR and of PolA to the *Bacillus subtilis* replication forks. Nucleic Acids Research. 2019 Sep 19;47(16):8521–36. doi:10.1093/nar/gkz554

40. Grimm JB, Xie L, Casler JC, Patel R, Tkachuk AN, Falco N, et al. A General Method to Improve Fluorophores Using Deuterated Auxochromes. JACS Au. 2021 May 24;1(5):690–6. doi:10.1021/jacsau.1c00006

41. Soubry N, Wang A, Reyes-Lamothe R. Replisome activity slowdown after exposure to ultraviolet light in *Escherichia coli*. Proc Natl Acad Sci USA. 2019 Jun 11;116(24):11747–53. doi:10.1073/pnas.1819297116

42. Uphoff S. Real-time dynamics of mutagenesis reveal the chronology of DNA repair and damage tolerance responses in single cells. Proc Natl Acad Sci USA. 2018 Jul 10;115(28). doi:10.1073/pnas.1801101115

43. Hernández-Tamayo R, Graumann PL. Bacillus subtilis RarA forms damage-inducible foci that scan the entire cell. BMC Res Notes. 2019 Dec;12(1):219. doi:10.1186/s13104-019-4252-x

44. Mallik S, Popodi EM, Hanson AJ, Foster PL. Interactions and Localization of Escherichia coli Error-Prone DNA Polymerase IV after DNA Damage. Gourse RL, editor. J Bacteriol. 2015 Sep;197(17):2792–809. doi:10.1128/JB.00101-15

45. Klocko AD, Schroeder JW, Walsh BW, Lenhart JS, Evans ML, Simmons LA. Mismatch repair causes the dynamic release of an essential DNA polymerase from the replication fork: Mismatch repair releases DnaE. Molecular Microbiology. 2011 Nov;82(3):648–63. doi:10.1111/j.1365-2958.2011.07841.x

46. Hernández-Tamayo R, Schmitz H, Graumann PL. Single-Molecule Dynamics at a Bacterial Replication Fork after Nutritional Downshift or Chemically Induced Block in Replication. Bowman GR, editor. mSphere. 2021 Feb 24;6(1):e00948–20. doi:10.1128/mSphere.00948-20

47. Zawadzki P, Stracy M, Ginda K, Zawadzka K, Lesterlin C, Kapanidis AN, et al. The Localization and Action of Topoisomerase IV in *Escherichia coli* Chromosome Segregation Is Coordinated by the SMC Complex, MukBEF. Cell Reports. 2015 Dec;13(11):2587–96. doi:10.1016/j.celrep.2015.11.034

48. Uphoff S, Reyes-Lamothe R, Garza De Leon F, Sherratt DJ, Kapanidis AN. Single-molecule DNA repair in live bacteria. Proc Natl Acad Sci USA. 2013 May 14;110(20):8063–8. doi:10.1073/pnas.1301804110

49. Lepore A, Thédié D, McLaren L, Goossens L, Azeroglu B, Pambos OJ, et al. *In vivo* single-molecule imaging of RecB reveals efficient repair of DNA damage in *Escherichia coli*. Nucleic Acids Research. 2025 May 22;53(10):gkaf454. doi:10.1093/nar/gkaf454

50. Becherel OJ, Fuchs RPP, Wagner J. Pivotal role of the β-clamp in translesion DNA synthesis and mutagenesis in E. coli cells. DNA Repair (Amst). 2002 Sep 4;1(9):703–8. doi:10.1016/s1568-7864(02)00106-4 PubMed PMID: 12509274.

51. Costes A, Lecointe F, McGovern S, Quevillon-Cheruel S, Polard P. The C-Terminal Domain of the Bacterial SSB Protein Acts as a DNA Maintenance Hub at Active Chromosome Replication Forks. Matic I, editor. PLoS Genet. 2010 Dec 9;6(12):e1001238. doi:10.1371/journal.pgen.1001238

52. Zeigler DR, Prágai Z, Rodriguez S, Chevreux B, Muffler A, Albert T, et al. The Origins of 168, W23, and Other *Bacillus subtilis* Legacy Strains. J Bacteriol. 2008 Nov;190(21):6983–95. doi:10.1128/JB.00722-08

53. Schroeder JW, Simmons LA. Complete Genome Sequence of *Bacillus subtilis* Strain PY79. Genome Announc. 2013 Dec 26;1(6):e01085–13. doi:10.1128/genomeA.01085-13

54. Tokunaga M, Imamoto N, Sakata-Sogawa K. Highly inclined thin illumination enables clear single-molecule imaging in cells. Nat Methods. 2008 Feb;5(2):159–61. doi:10.1038/nmeth1171

55. Sliusarenko O, Heinritz J, Emonet T, Jacobs-Wagner C. High-throughput, subpixel precision analysis of bacterial morphogenesis and intracellular spatio-temporal dynamics. Molecular Microbiology. 2011 May;80(3):612–27. doi:10.1111/j.1365-2958.2011.07579.x

56. Jaqaman K, Loerke D, Mettlen M, Kuwata H, Grinstein S, Schmid SL, et al. Robust single-particle tracking in live-cell time-lapse sequences. Nat Methods. 2008 Aug;5(8):695–702. doi:10.1038/nmeth.1237

57. Aguet F, Antonescu CN, Mettlen M, Schmid SL, Danuser G. Advances in Analysis of Low Signal-to-Noise Images Link Dynamin and AP2 to the Functions of an Endocytic Checkpoint. Developmental Cell. 2013 Aug;26(3):279–91. doi:10.1016/j.devcel.2013.06.019

