## Supplementary Information for "“The *B. subtilis* translesion polymerase Pol Y1 is not strongly recruited to sites of replication upon different types of DNA damage”"

**This PDF file includes:**

Figures S1 to S3
Tables S1 to S8
Supplementary Methods
Supplementary References

**Supplementary Figures**

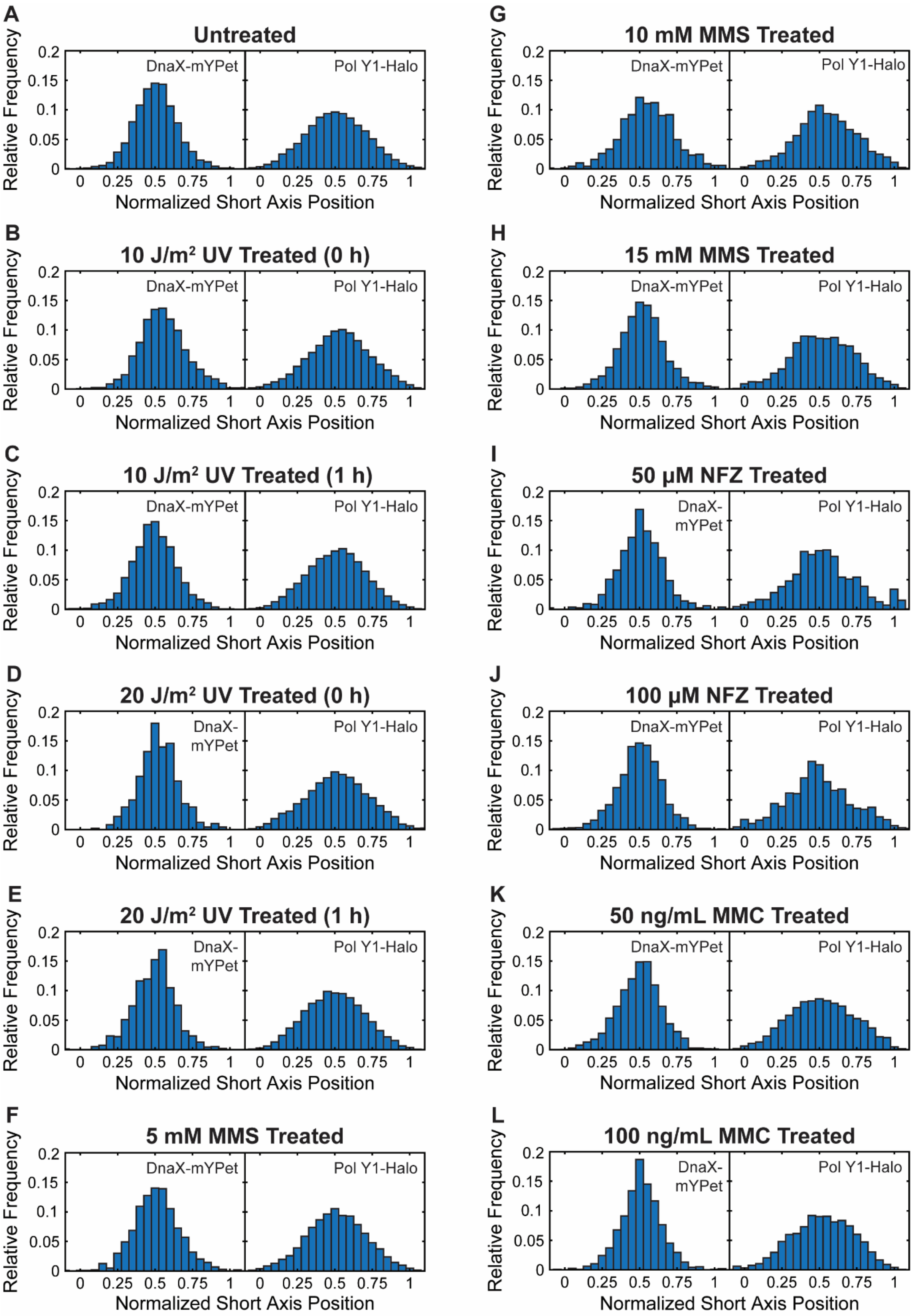

A
Untreated
B
10 J/m² UV Treated (0 h)
C
10 J/m² UV Treated (1 h)
D
20 J/m² UV Treated (0 h)
E
20 J/m² UV Treated (1 h)
F
5 mM MMS Treated
G
10 mM MMS Treated
H
15 mM MMS Treated
I
50 μM NFZ Treated
J
100 μM NFZ Treated
K
50 ng/mL MMC Treated
L
100 ng/mL MMC Treated
DnaX-mYPet
Pol Y1-Halo
Relative Frequency
0
0.05
0.1
0.15
0.2
0
0.25
0.5
0.75
1
Normalized Short Axis Position

**Figure S1.** Average cellular localization of DnaX-mYPet and Pol Y1-Halo in the presence of different types of DNA damage. Short axis projections of DnaX foci (left) and Pol Y1 trajectories (right) in (A) untreated cells and after treatment with (B) 10 J/m$^2$ 254 nm UV light ($t$ = 0 h), (C) 10 J/m$^2$ 254 nm UV light ($t$ = 1 h), (D) 20 J/m$^2$ 254 nm UV light ($t$ = 0 h), (E) 20 J/m$^2$ 254 nm UV light ($t$ = 1 h), (F) 5 mM MMS, (G) 10 mM MMS, (H) 15 mM MMS, (I) 50 µM NFZ, (J) 100 µM NFZ, (K) 50 ng/mL MMC, and (L) 100 ng/mL MMC.

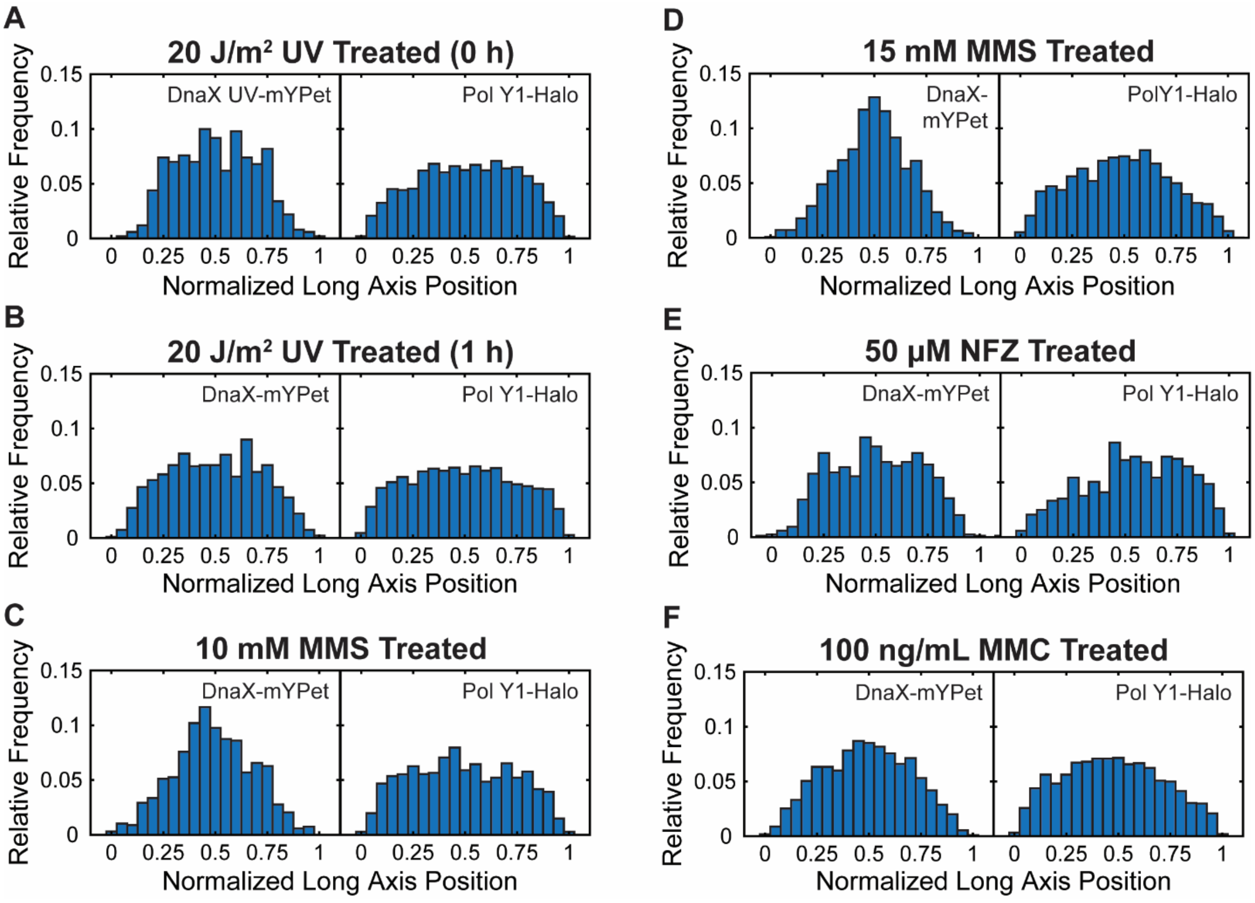


**Figure S2.** Average cellular localization of DnaX-mYPet and Pol Y1-Halo in the presence of different types of DNA damage. Long axis projections of DnaX foci (left) and Pol Y1 trajectories (right) after treatment with (A) 20 J/m$^2$ 254 nm UV light ($t$ = 0 h), (B) 20 J/m$^2$ 254 nm UV light ($t$ = 1 h), (C) 10 mM MMS, (D) 15 mM MMS, (E) 50 µM NFZ, and (F) 100 ng/mL MMC.

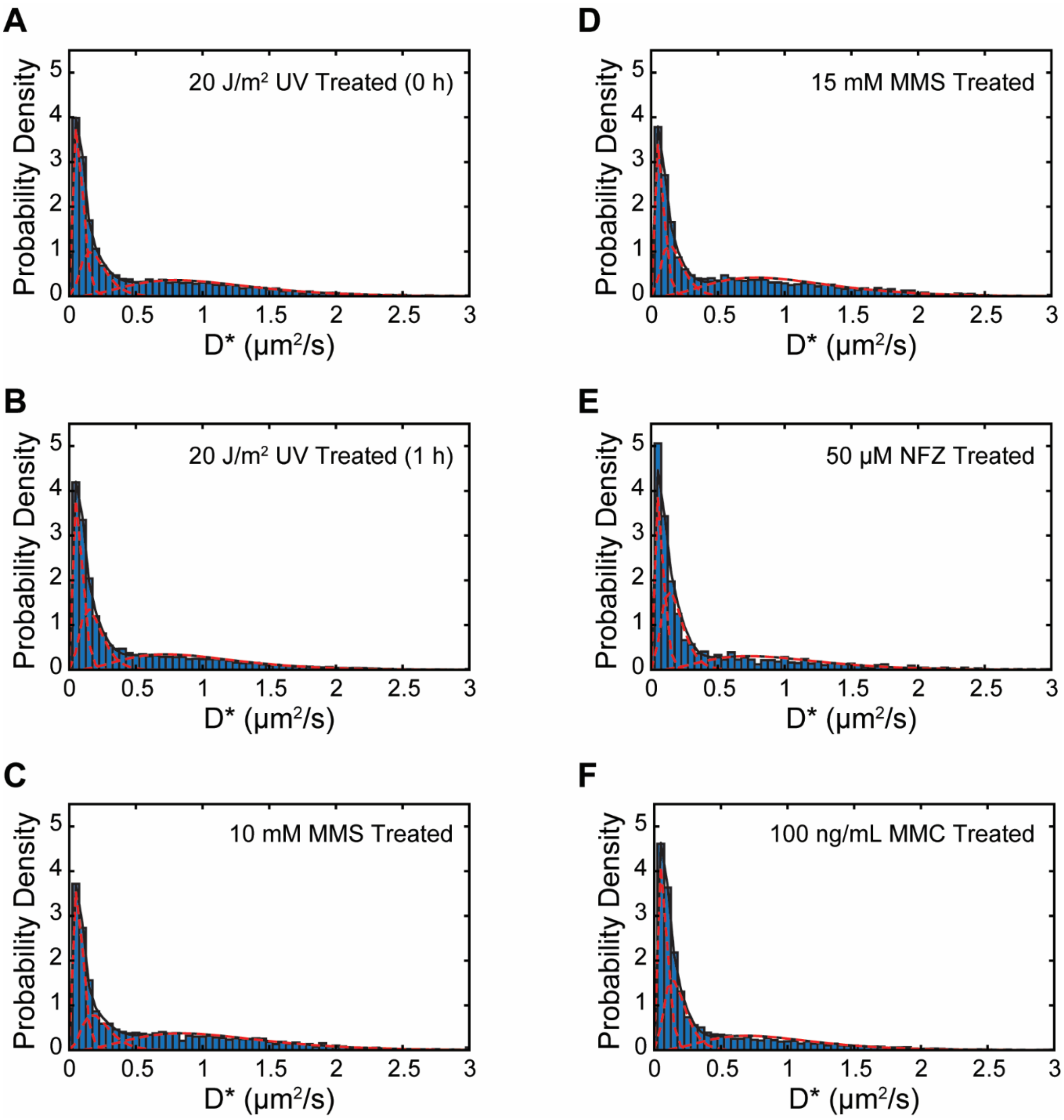


**Figure S3.** Apparent diffusion coefficient ($D^*$) distributions and corresponding three-population fits for Pol Y1-Halo in the presence of different types of DNA damage. Pol Y1 $D^*$ distributions in after treatment with (A) 20 J/m$^2$ 254 nm UV light ($t$ = 0 h), (B) 20 J/m$^2$ 254 nm UV light ($t$ = 1 h), (C) 10 mM MMS, (D) 15 mM MMS, (E) 50 µM NFZ, and (F) 100 ng/mL MMC. Individual populations are shown as red dashed lines and overall fits are shown as solid black lines.

### Supplementary Tables

**Table S1.** Mutagenesis for different strains under different treatment conditions measured by the rate of rifampicin resistance ($Rif^R$) (mean ± standard deviation). All MMS, NFZ, and MMC incubations were performed for 1 h. A separate untreated dataset was collected in matched experiments for each treatment condition. (Note: * indicates statistically significant difference at the $p < 0.05$ level relative to WT for same treatment condition. ‡ indicates statistically significant difference at the $p < 0.05$ level for treated vs. untreated condition for the same strain.)

| **Strain** | **WT** | | ***ΔPol Y1*** | | ***ΔPol Y2*** | | ***ΔPol Y1 ΔPol Y2*** | |
|---|---|---|---|---|---|---|---|---|
| **Condition** | **Untreated** | **40 J/m$^2$ UV** | **Untreated** | **40 J/m$^2$ UV** | **Untreated** | **40 J/m$^2$ UV** | **Untreated** | **40 J/m$^2$ UV** |
| **$Rif^R$ (per $10^8$)** | 0.9 ± 0.6 | 16 ± 4 *‡ | 0.7 ± 0.2 | 5 ± 3 *‡ | 0.6 ± 0.3 | 5 ± 1 *‡ | 0.7 ± 0.4 | 0.6 ± 0.6 * |
| **Condition** | **Untreated** | **10 mM MMS** | **Untreated** | **10 mM MMS** | **Untreated** | **10 mM MMS** | **Untreated** | **10 mM MMS** |
| **$Rif^R$ (per $10^8$)** | 1.5 ± 0.8 | 11 ± 2 ‡ | 2 ± 1 | 9 ± 5 ‡ | 0.8 ± 0.6 | 10 ± 5 ‡ | 1 ± 1 | 8 ± 3 ‡ |
| **Condition** | **Untreated** | **100 µM NFZ** | **Untreated** | **100 µM NFZ** | **Untreated** | **100 µM NFZ** | **Untreated** | **100 µM NFZ** |
| **$Rif^R$ (per $10^8$)** | 1.5 ± 0.3 | 3.2 ± 0.6 ‡ | 1.9 ± 0.6 | 4 ± 1 ‡ | 1.4 ± 0.6 | 3 ± 2 | 1.8 ± 0.7 | 3 ± 1 |
| **Condition** | **Untreated** | **200 ng/mL MMC** | **Untreated** | **200 ng/mL MMC** | **Untreated** | **200 ng/mL MMC** | **Untreated** | **200 ng/mL MMC** |
| **$Rif^R$ (per $10^8$)** | 0.8 ± 0.6 | 12 ± 5 ‡ | 0.8 ± 0.5 | 14 ± 7 ‡ | 1.5 ± 0.9 | 2.1 ± 0.6 * | 1.3 ± 0.5 | 2 ± 1 * |

**Table S2.** Cell length and width and number of DnaX foci per cell (mean ± standard error of the mean (S.E.M.)) under different treatment conditions. (Note: ‡ indicates difference was not statistically significant at the $p < 0.05$ level relative to the untreated condition.)

| **Strain** | **Cell Length (µm)** | **Cell Width (µm)** | **Number of Foci per Cell** |
|---|---|---|---|
| Untreated | 3.40 ± 0.02 | 0.699 ± 0.002 | 1.76 ± 0.02 |
| 10 J/m$^2$ UV (0 h) | 3.40 ± 0.03 ‡ | 0.680 ± 0.002 | 1.67 ± 0.03 |
| 10 J/m$^2$ UV (1 h) | 4.07 ± 0.04 | 0.698 ± 0.002 ‡ | 2.13 ± 0.04 |
| 20 J/m$^2$ UV (0 h) | 3.64 ± 0.05 | 0.708 ± 0.004 ‡ | 1.70 ± 0.05 ‡ |
| 20 J/m$^2$ UV (1 h) | 4.10 ± 0.04 | 0.692 ± 0.003 ‡ | 2.13 ± 0.05 |
| 5 mM MMS (1 h) | 3.19 ± 0.03 | 0.711 ± 0.002 | 1.74 ± 0.04 ‡ |
| 10 mM MMS (1 h) | 3.15 ± 0.04 | 0.664 ± 0.002 | 1.41 ± 0.04 |
| 15 mM MMS (1 h) | 3.14 ± 0.03 | 0.677 ± 0.002 | 1.50 ± 0.02 |
| 50 µM NFZ (1 h) | 2.95 ± 0.04 | 0.621 ± 0.003 | 1.46 ± 0.04 |
| 100 µM NFZ (1 h) | 2.72 ± 0.02 | 0.616 ± 0.002 | 1.31 ± 0.02 |
| 50 ng/mL MMC (1 h) | 3.99 ± 0.03 | 0.672 ± 0.002 | 1.83 ± 0.03 ‡ |
| 100 ng/mL MMC (1 h) | 4.07 ± 0.04 | 0.681 ± 0.003 | 1.74 ± 0.04 ‡ |

**Table S3.** Fold change in number of colony forming units per mL (CFUs/mL) for imaging cultures after different treatments (mean ± standard deviation).

| **Condition** | **Untreated** | **DMF (1 h)** | **DMSO (1 h)** | |
|---|---|---|---|---|
| **Fold Change in CFUs/mL** | 2.1 ± 0.4 | 1.9 ± 0.3 | 1.9 ± 0.2 | |
| **Condition** | **10 J/m$^2$ UV (0 h)** | **10 J/m$^2$ UV (1 h)** | **20 J/m$^2$ UV (0 h)** | **20 J/m$^2$ UV (1 h)** |
| **Fold Change in CFUs/mL** | 0.72 ± 0.08 | 1.3 ± 0.2 | 0.4 ± 0.1 | 0.9 ± 0.2 |
| **Condition** | **5 mM MMS (1 h)** | **10 mM MMS (1 h)** | **15 mM MMS (1 h)** | |
| **Fold Change in CFUs/mL** | 1.4 ± 0.3 | 0.9 ± 0.1 | 0.30 ± 0.03 | |
| **Condition** | **50 µM NFZ (1 h)** | **100 µM NFZ (1 h)** | **250 µM NFZ (1 h)** | |
| **Fold Change in CFUs/mL** | 1.1 ± 0.3 | 1.0 ± 0.1 | 1.2 ± 0.2 | |
| **Condition** | **50 ng/mL MMC (1 h)** | **100 ng/mL MMC (1 h)** | | |
| **Fold Change in CFUs/mL** | 0.8 ± 0.2 | 0.5 ± 0.1 | | |

**Table S4.** Value of the mean radial distribution function $g(r)$ for Pol Y1-DnaX colocalization at the second smallest value of $r$ (generally the maximum of the $g(r)$ curve) and the standard error of the mean (S.E.M.) at that $r$ value for the 100 calculated $g(r)$ curves.

| **Figure(s)** | **Condition** | **$g(r)$ ± S.E.M.** |
|---|---|---|
| 4A, 4B, 4C, 4D | Untreated | 2.000 ± 0.007 |
| 4A | 10 J/m$^2$ UV (0 h) | 2.214 ± 0.009 |
| | 10 J/m$^2$ UV (1 h) | 2.371 ± 0.009 |
| | 20 J/m$^2$ UV (0 h) | 2.59 ± 0.05 |
| | 20 J/m$^2$ UV (1 h) | 2.69 ± 0.03 |
| 4B | 5 mM MMS (1 h) | 2.03 ± 0.05 |
| | 10 mM MMS (1 h) | 2.64 ± 0.05 |
| | 15 mM MMS (1 h) | 2.38 ± 0.03 |
| 4C | 50 µM NFZ (1 h) | 4.46 ± 0.08 |
| | 100 µM NFZ (1 h) | 2.62 ± 0.03 |
| 4D | 50 ng/mL MMC (1 h) | 2.82 ± 0.03 |
| | 100 ng/mL MMC (1 h) | 2.74 ± 0.06 |

**Table S5.** Halo-DnaE and PolC-Halo diffusion coefficient distribution fit parameters from MSD analysis (± uncertainties from 95% fit confidence intervals).

| Condition | $D_1$ (μm²/s) | $A_1$ | $D_2$ (μm²/s) | $A_2$ | $D_3$ (μm²/s) | $A_3$ |
|---|---|---|---|---|---|---|
| Untreated | 0.082 ± 0.004 | 0.317 ± 0.032 | 1.0761 ± 0.086 | 0.477 ± 0.028 | 0.230 ± 0.036 | 0.206 ± 0.060 |
| 10 J/m² UV (0 h) | 0.077 ± 0.005 | 0.299 ± 0.033 | 1.065 ± 0.087 | 0.473 ± 0.028 | 0.214 ± 0.031 | 0.228 ± 0.061 |
| 10 J/m² UV (1 h) | 0.078 ± 0.083 | 0.324 ± 0.034 | 1.025 ± 0.095 | 0.439 ± 0.030 | 0.221 ± 0.032 | 0.240 ± 0.065 |
| 20 J/m² UV (0 h) | 0.079 ± 0.005 | 0.343 ± 0.039 | 1.065 ± 0.099 | 0.426 ± 0.029 | 0.209 ± 0.032 | 0.231 ± 0.067 |
| 20 J/m² UV (1 h) | 0.077 ± 0.005 | 0.336 ± 0.045 | 0.972 ± 0.098 | 0.376 ± 0.027 | 0.190 ± 0.025 | 0.289 ± 0.075 |
| 5 mM MMS (1 h) | 0.081 ± 0.004 | 0.288 ± 0.025 | 1.134 ± 0.07 | 0.531 ± 0.023 | 0.230 ± 0.033 | 0.819 ± 0.048 |
| 10 mM MMS (1 h) | 0.079 ± 0.004 | 0.327 ± 0.031 | 1.123 ± 0.10 | 0.468 ± 0.030 | 0.231 ± 0.037 | 0.206 ± 0.061 |
| 15 mM MMS (1 h) | 0.073 ± 0.007 | 0.278 ± 0.052 | 1.051 ± 0.089 | 0.489 ± 0.027 | 0.180 ± 0.030 | 0.233 ± 0.080 |
| 50 μM NFZ (1 h) | 0.070 ± 0.007 | 0.358 ± 0.062 | 1.002 ± 0.157 | 0.334 ± 0.035 | 0.179 ± 0.029 | 0.308 ± 0.097 |
| 100 μM NFZ (1 h) | 0.074 ± 0.005 | 0.329 ± 0.038 | 0.959 ± 0.103 | 0.419 ± 0.034 | 0.212 ± 0.033 | 0.252 ± 0.072 |
| 50 ng/mL MMC (1 h) | 0.076 ± 0.005 | 0.437 ± 0.056 | 1.005 ± 0.156 | 0.325 ± 0.035 | 0.195 ± 0.039 | 0.238 ± 0.092 |
| 100 ng/mL MMC (1 h) | 0.075 ± 0.007 | 0.351 ± 0.064 | 0.931 ± 0.125 | 0.332 ± 0.032 | 0.180 ± 0.027 | 0.317 ± 0.096 |

**Table S6.** Oligonucleotides used in this study. (Lowercase letters indicate bases that do not prime on the template but provide homology for Gibson assembly of PCR fragments.)

| Number | Designation | Sequence (5′-3′) |
|---|---|---|
| oEST028 | yqjH-downstream-rev | CGACAACTTCAGATGGGCTGGTGTTC |
| oEST037 | yqjH-upstream-for | CATCAGTCACCGTATTGACT |
| oEST038 | yqjH-Nter-rev | TCGGCTCTTTCCCGGCATAA |
| oEST039 | yqjH-Nter-spec-iso-for | ttatgccgggaaagagccgaGGATCCTTCTGCTCCCTCGC |
| oEST040 | yqjH-Cter-spec-iso-rev | attcagcttttcttttcatcGACCAGGGAGCACTGGTCAA |
| oEST041 | yqjH-Cter-for | GATGAAAAGAAAAGCTGAATCGC |

**Table S7.** *B. subtilis* bacterial strains used in this study.

| Number | Designation or description | Relevant genotype | Construction or source strain designation | Reference |
|---|---|---|---|---|
| EST003 (WT) | *B. subtilis* prototrophic wild-type strain | PY79 | — | (1, 2) |
| EST0081 | PolC-Dendra2 | PY79 *polC-dendra2 loxP-spec-loxP* | Gift of Xindan Wang (Indiana University); strain BWX2913 | — |
| EST111 | ΔPol Y1 | PY79 *yqjH::loxP-spec-loxP* | EST111 | (3) |
| EST117 | ΔPol Y2 | PY79 *yqjW::loxP-spec-loxP* | EST117 | (3) |
| EST137 | ΔPol Y1 | PY79 *yqjH::loxP-kan-loxP* | Transformation: *yqjH::loxP-kan-loxP* → EST003 | This study |
| EST169 | ΔPol Y1 ΔPol Y2 | PY79 *yqjH::loxP-kan-loxP yqjW::loxP-spec-loxP* | Transformation: EST117 → EST137 | This study |
| EST197 | Pol Y1-Halo DnaX-mYPet | PY79 *yqjH-halo loxP spec dnaX-mYpet cat Ω pWX340a* | EST197 | (3) |

**Table S8.** Imaging dataset size.

| Dataset/Condition | Figure(s) | Number of Days | Number of Replicates | Number of Cells | Number of Tracks or Foci |
|---|---|---|---|---|---|
| DnaX cellular localization Untreated | 3C, S1A | 6 | 23 | 1,881 | 3,316 |
| Pol Y1 cellular localization Untreated | 3C, S1A | 6 | 23 | 1,881 | 31,649 |
| DnaX cellular localization 10 J/m$^2$ UV (0 h) | 3D, S1B | 4 | 14 | 909 | 1,520 |
| Pol Y1 cellular localization 10 J/m$^2$ UV (0 h) | 3D, S1B | 4 | 14 | 909 | 18,925 |
| DnaX cellular localization 10 J/m$^2$ UV (1 h) | 3E, S1C | 4 | 14 | 858 | 1,828 |
| Pol Y1 cellular localization 10 J/m$^2$ UV (1 h) | 3E, S1C | 4 | 14 | 858 | 27,667 |
| DnaX cellular localization 20 J/m$^2$ UV (0 h) | 3F, S1D | 3 | 9 | 296 | 501 |
| Pol Y1 cellular localization 20 J/m$^2$ UV (0 h) | 3F, S1D | 3 | 9 | 296 | 7,519 |
| DnaX cellular localization 20 J/m$^2$ UV (1 h) | 3G, S1E | 3 | 10 | 445 | 946 |
| Pol Y1 cellular localization 20 J/m$^2$ UV (1 h) | 3G, S1E | 3 | 10 | 445 | 14,993 |
| DnaX cellular localization | 3H, S1F | 3 | 10 | 594 | 1,034 |

| | | | | | |
|---|---|---|---|---|---|
| 5 mM MMS (1 h) | | | | | |
| Pol Y1 cellular localization<br>5 mM MMS (1 h) | 3H, S1F | 3 | 10 | 594 | 7,023 |
| DnaX cellular localization<br>10 mM MMS (1 h) | 3I, S1G | 3 | 9 | 488 | 686 |
| Pol Y1 cellular localization<br>10 mM MMS (1 h) | 3I, S1G | 3 | 9 | 488 | 4,108 |
| DnaX cellular localization<br>15 mM MMS (1 h) | 3J, S1H | 4 | 13 | 945 | 1,410 |
| Pol Y1 cellular localization<br>15 mM MMS (1 h) | 3J, S1H | 4 | 13 | 945 | 4,900 |
| DnaX cellular localization<br>50 µM NFZ (1 h) | 3K, S1I | 4 | 12 | 579 | 846 |
| Pol Y1 cellular localization<br>50 µM NFZ (1 h) | 3K, S1I | 4 | 12 | 579 | 1,565 |
| DnaX cellular localization<br>100 µM NFZ (1 h) | 3L, S1J | 5 | 18 | 1,638 | 2,148 |
| Pol Y1 cellular localization<br>100 µM NFZ (1 h) | 3L, S1J | 5 | 18 | 1,638 | 3,379 |
| DnaX cellular localization<br>50 ng/mL MMC (1 h) | 3M, S1K | 4 | 13 | 981 | 1,800 |
| Pol Y1 cellular localization<br>50 ng/mL MMC (1 h) | 3M, S1K | 4 | 13 | 981 | 10,325 |
| DnaX cellular localization<br>100 ng/mL MMC (1 h) | 3N, S1L | 4 | 12 | 663 | 1,151 |
| Pol Y1 cellular localization<br>100s ng/mL MMC (1 h) | 3N, S1L | 4 | 12 | 663 | 8,235 |
| Pol Y1-DnaX *g*(*r*)<br>Untreated | 4A, 4B,<br>4C, 4D | 6 | 23 | 1,881 | 30,606 |
| Pol Y1-DnaX *g*(*r*)<br>10 J/m$^2$ UV (0 h) | 4A | 4 | 14 | 909 | 18,441 |
| Pol Y1-DnaX *g*(*r*)<br>10 J/m$^2$ UV (1 h) | 4A | 4 | 14 | 858 | 27,173 |
| Pol Y1-DnaX *g*(*r*)<br>20 J/m$^2$ UV (0 h) | 4A | 3 | 9 | 296 | 2,482 |
| Pol Y1-DnaX *g*(*r*)<br>20 J/m$^2$ UV (1 h) | 4A | 3 | 10 | 445 | 5,864 |
| Pol Y1-DnaX *g*(*r*)<br>5 mM MMS (1 h) | 4B | 3 | 10 | 594 | 1,614 |
| Pol Y1-DnaX *g*(*r*)<br>10 mM MMS (1 h) | 4B | 3 | 9 | 488 | 1,878 |
| Pol Y1-DnaX *g*(*r*)<br>15 mM MMS (1 h) | 4B | 4 | 13 | 945 | 4,722 |
| Pol Y1-DnaX *g*(*r*)<br>50 µM NFZ (1 h) | 4C | 4 | 12 | 579 | 1,454 |
| Pol Y1-DnaX *g*(*r*)<br>100 µM NFZ (1 h) | 4C | 5 | 18 | 1,638 | 3,033 |
| Pol Y1-DnaX *g*(*r*)<br>50 ng/mL MMC (1 h) | 4D | 4 | 13 | 981 | 3,323 |
| Pol Y1-DnaX *g*(*r*)<br>100 ng/mL MMC (1 h) | 4D | 4 | 12 | 663 | 1,913 |
| Pol Y1 $D^*$<br>Untreated | 5A | 6 | 23 | 1,881 | 29,120 |
| Pol Y1 $D^*$<br>10 J/m$^2$ UV (0 h) | 5B | 4 | 14 | 909 | 17,446 |

| | | | | | |
|---|---|---|---|---|---|
| Pol Y1 *D**<br>10 J/m$^2$ UV (1 h) | 5C | 4 | 14 | 858 | 25,449 |
| Pol Y1 *D**<br>20 J/m$^2$ UV (0 h) | 5D | 3 | 9 | 296 | 6,881 |
| Pol Y1 *D**<br>20 J/m$^2$ UV (1 h) | 5E | 3 | 10 | 445 | 13,760 |
| Pol Y1 *D**<br>5 mM MMS (1 h) | 5F | 3 | 10 | 594 | 6,462 |
| Pol Y1 *D**<br>10 mM MMS (1 h) | 5G | 3 | 9 | 488 | 3,729 |
| Pol Y1 *D**<br>15 mM MMS (1 h) | 5H | 4 | 13 | 945 | 4,512 |
| Pol Y1 *D**<br>50 μM NFZ (1 h) | 5I | 4 | 12 | 579 | 1,390 |
| Pol Y1 *D**<br>100 μM NFZ (1 h) | 5J | 5 | 18 | 1,638 | 2,993 |
| Pol Y1 *D**<br>50 ng/mL MMC (1 h) | 5K | 4 | 13 | 981 | 9,385 |
| Pol Y1 *D**<br>100 ng/mL MMC (1 h) | 5L | 4 | 12 | 663 | 7,381 |

### Supplementary Methods

*Overview of strain construction strategy:*

Introduction of chromosomal genetic modifications by transformation of genomic DNA or double-stranded DNA (dsDNA) fragments assembled via Gibson assembly(4) was performed as described previously.(3) All oligonucleotides and bacterial strains used in this study are listed in Tables S1 and S2, respectively. Construction details for all new strains are summarized below.

*Detailed strain construction information:*

**EST137:** ΔPol Y1. The *yqjH* upstream was amplified from strain EST003 using oligonucleotides oEST037 and oEST038. The loxP spec cassette was amplified from strain EST081 using oligonucleotides oEST039 and oEST040. The *yqjH* downstream was amplified from strain EST003 using oligonucleotides oEST041 and oEST028. The three fragments were joined by Gibson assembly and transformed into strain EST003.

**EST 169:** ΔPol Y1 ΔPol Y2. The *yqjW::loxP-spec-loxP* allele was transferred from strain EST117 to strain EST137 by transformation with genomic DNA.

### Supplementary References


1. Zeigler,D.R., Prágai,Z., Rodriguez,S., Chevreux,B., Muffler,A., Albert,T., Bai,R., Wyss,M. and Perkins,J.B. (2008) The Origins of 168, W23, and Other *Bacillus subtilis* Legacy Strains. *J Bacteriol*, **190**, 6983–6995.

2. Schroeder,J.W. and Simmons,L.A. (2013) Complete Genome Sequence of *Bacillus subtilis* Strain PY79. *Genome Announc*, **1**, e01085-13.

3. Marrin,M.E., Foster,M.R., Santana,C.M., Choi,Y., Jassal,A.S., Rancic,S.J., Greenwald,C.R., Drucker,M.N., Feldman,D.T. and Thrall,E.S. (2024) The translesion polymerase Pol Y1 is a constitutive component of the *B. subtilis* replication machinery. *Nucleic Acids Research*, **52**, 9613–9629.

4. Gibson,D.G., Young,L., Chuang,R.-Y., Venter,J.C., Hutchison,C.A. and Smith,H.O. (2009) Enzymatic assembly of DNA molecules up to several hundred kilobases. *Nat Methods*, **6**, 343–345.
